# T cells discriminate between groups C1 and C2 HLA-C

**DOI:** 10.1101/2021.11.11.468262

**Authors:** Malcolm J. W. Sim, Zachary Stotz, Jinghua Lu, Paul Brennan, Eric O. Long, Peter D. Sun

## Abstract

Dimorphic residues at positions 77 and 80 delineate HLA-C allotypes into two groups, C1 and C2, which associate with disease through interactions with C1 and C2-specific natural killer cell receptors. How the C1/C2 dimorphism affects T cell recognition is unknown. Using HLA-C allotypes that differ only by the C1/C2-defining residues, we found that KRAS-G12D neoantigen specific T cell receptors (TCR) discriminated groups C1 and C2 HLA-C, due to effects on peptide presentation and TCR affinity. Structural and functional experiments combined with immunopeptidomics analysis revealed that C1-HLA-C favors smaller amino acids at the peptide C-terminus minus-1 position (pΩ-1), and that larger pΩ-1 residues diminished TCR recognition of C1-HLA-C. After controlling for peptide presentation, TCRs exhibited weaker affinities for C2-HLA-C despite conserved TCR contacts. Thus, the C1/C2 dimorphism impacts peptide presentation and HLA-C restricted T cell responses, with implications in multiple disease contexts including adoptive T cell therapy targeting KRAS-G12D-induced cancers.

## Introduction

The classical class I human leukocyte antigens (HLA-I; HLA-A, HLA-B and HLA-C) are the most polymorphic genes across human populations and are associated with a myriad of human diseases (Dendrou et al., 2018; Robinson et al., 2017). HLA-I molecules present short peptides on the cell surface of nucleated cells where they form ligands for receptors of the immune system including T cell receptors (TCR) and natural killer (NK) cell receptors (Rossjohn et al., 2015; Saunders et al., 2015). HLA-I bound peptides are collectively known as the immunopeptidome, which defines the subset of the proteome bound as peptides to a particular HLA-I allotype (Istrail et al., 2004). Immunopeptidomes can be highly diverse, consisting of hundreds to thousands of different sequences as it is shaped by polymorphic amino acid residues that form pockets (A to F) in the HLA-I peptide binding groove (PBG) (Di Marco et al., 2017; Madden, 1995; Sarkizova et al., 2020). These pockets restrict the peptide repertoire to those peptides with certain amino acid side chains at particular positions, which form anchor residues that define allotype specific motifs (Di Marco et al., 2017; Falk et al., 1991; Sarkizova et al., 2020). The ‘anchor’ residues are often position 2 (p2) or p3 (B and C pockets) and the C-terminus (pΩ; F pocket), while residues outside the anchors can vary extensively allowing immunopeptidome diversification (Di Marco et al., 2017; Falk et al., 1991; Madden, 1995; Sarkizova et al., 2020). HLA-I allotypes with similar PBGs bind similar peptides but subtle amino acid differences can modify the side chain orientation impacting interactions with TCRs and NK cell receptors (Illing et al., 2018; Illing et al., 2012; Saunders et al., 2020; Stewart-Jones et al., 2012; Tynan et al., 2005). Understanding how HLA-I polymorphism impacts the immunopeptidome and interactions with immunoreceptors is critical for constructing molecular mechanisms that underpin disease risk associated with specific HLA-I alleles (Dendrou et al., 2018).

HLA-C is thought to play a lesser role in T cell responses compared to that of HLA-A and HLA-B due to its lower cell surface expression level (Apps et al., 2015; McCutcheon et al., 1995). However, HLA-C restricted T cells are implicated in several disease settings. In HIV infection, higher expressing HLA-C allotypes associated with increased T cell responses, while the converse association was found with Chron’s disease (Apps et al., 2013). In psoriasis, the strongest genetic association is with *HLA-C*06:02*, thought to be mediated by HLA-C*06:02 restricted T cells (Chen and Tsai, 2018; Lande et al., 2014). Furthermore, recent efforts to define the specificities of tumor infiltrating lymphocytes (TIL), readily identified HLA-C restricted T cells (Levin et al., 2021; Lu et al., 2014; Murata et al., 2020; Tran et al., 2015; Tran et al., 2016). Most notably, multiple HLA-C*08:02 restricted TCRs specific for the oncogenic hotspot mutation KRAS-G12D were identified in multiple cancer patients (Tran et al., 2015; Tran et al., 2016). Adoptive transfer of expanded KRAS-G12D specific TILs successfully treated a patient with metastatic colorectal cancer leading to complete regression of all but one metastatic lesion (Tran et al., 2016). The remaining lesion lost *HLA-C*08:02* from its genome demonstrating conclusively that HLA-C restricted T cells can mediate effective anti tumour responses (Tran et al., 2016). Despite the clinical relevance of HLA-C restricted T cells, few studies have examined TCR-HLA-C interactions with a molecular and structural focus. Indeed, our recent study on HLA-C*08:02 restricted KRAS-G12D specific TCRs, was the first to present crystal structures of any TCR in complex with HLA-C (Sim et al., 2020).

*HLA-C* is the most recently evolved classical HLA-I gene, present in only humans and other apes, and has several unique features (Adams and Parham, 2001). HLA-C, not HLA-A or HLA-B, is expressed on extra-villous trophoblasts, making it the most polymorphic molecule expressed at the maternal-fetal interface (King et al., 2000). In addition, all HLA-C variants are ligands for the killer-cell immunoglobulin-like receptors (KIR) in contrast to only a subset of HLA-A and HLA-B (Hilton and Parham, 2017; Moesta et al., 2008). The KIR are a family of activating and inhibitory receptors expressed primarily on NK cells (Hilton and Parham, 2017; Saunders et al., 2015). HLA-C allotypes form two groups, C1 and C2 based on two dimorphic amino acid residues at positions 77 and 80 (Biassoni et al., 1995; Parham et al., 2012). C1 HLA-C allotypes carry Ser77 and Asn80 (Fig. 1A) and are ligands for the inhibitory receptors KIR2DL2/3. The inhibitory receptor KIR2DL1 binds C2 HLA-C allotypes, which carry Asn77 and Lys80. The KIR discriminate C1 and C2 HLA-C largely via position 44 in the KIR and direct interactions with HLA-C position 80 (Boyington et al., 2000; Boyington and Sun, 2002; Fan et al., 2001). Genetic association studies link C1/C2 status and the presence and absence of specific KIR-HLA combinations with risk of multiple human diseases including cancer, infectious diseases, autoimmunity and disorders of pregnancy (Hiby et al., 2004; Khakoo et al., 2004; Kulkarni et al., 2008; Littera et al., 2016; Parham and Moffett, 2013; Rajagopalan and Long, 2005; Venstrom et al., 2012). Whether the C1/C2 dimorphism impacts HLA-C interactions with other immunoreceptors is unknown.

**Figure 1.**
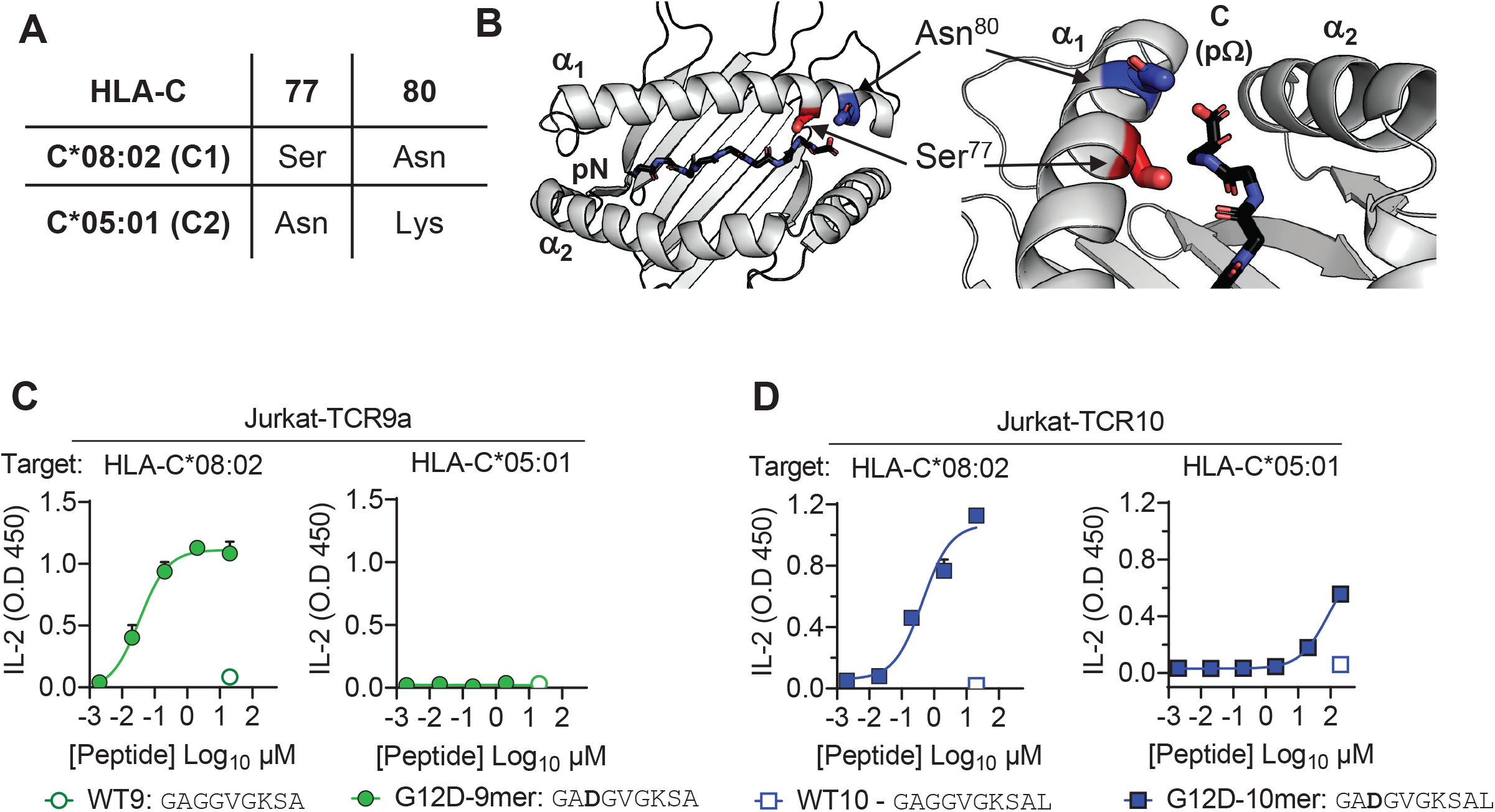
HLA-C C1/C2 dimorphism impacts T-cell recognition of KRAS-G12D neoantigen. **(A)** Sequence of C1/C2 dimorphism in HLA-C*08:02 and HLA-C*05:01 at positions 77 and 80. All other residues are conserved. **(B)** Location of C1/C2 dimorphism on structure of HLA-C*08:02 (PDB:6ULI). **(C-D)** Stimulation of TCR9a^+^ Jurkat cells **(C)** and TCR10^+^ Jurkat cells **(D)** by 221-C*08:02 (left) or 221-C*05:01 (right) preloaded with KRAS WT or G12D peptides at indicated concentrations. Concentration of IL-2 in culture supernatant was measure by ELISA. A representative experiment of three biological replicates is shown.

Given the clinical relevance of HLA-C restricted T cells and that all HLA-C allotypes are either C1 or C2, it is important to ask whether this dimorphism impacts T cell recognition. It was hitherto assumed that the C1/C2 dimorphism would not affect T cell recognition as it lies outside the general TCR footprint (Fig. 1A, B). To answer this question, we studied two HLA-C*08:02 (C1) restricted TCRs specific for different KRAS-G12D neoantigens. TCR9a is specific for G12D-9mer (GA**D**GVGKSA), while TCR10 is specific for G12D-10mer (GA**D**GVGKSAL) (Sim et al., 2020; Tran et al., 2016). TCR9a and TCR10 do not share TCR V genes and recognize their peptide antigens in different ways (Sim et al., 2020; Tran et al., 2016). We compared TCR9a and TCR10 recognition of HLA-C*08:02 (C1) with that of HLA-C*05:01, a C2 allotype that is identical in sequence to HLA-C*08:02 apart from the C1/C2 defining residues at positions 77 and 80 (Sim et al., 2017). In a previous study, we showed that 28 different ‘self’ peptides eluted from HLA-C*05:01 bound to both HLA-C*05:01 (C2) and HLA-C*08:02 (C1), suggesting a minimal impact of the C1/C2 dimorphism on peptide binding to HLA-C (Sim et al., 2017). Unexpectedly, we found that T cells discriminate between C1 and C2 HLA-C through impacts on both HLA-C peptide presentation and TCR binding, revealing unknown features of the C1/C2 dimorphism governing interactions of HLA-C with immunoreceptors. Further, our data suggest HLA-C*05:01 positive individuals may be eligible for immunotherapies targeting KRAS-G12D-10mer.

## Results

### Impact of the C1/C2 dimorphism to T-cell recognition of KRAS-neoantigens

HLA-C*05:01 (C*05) and HLA-C*08:02 (C*08) differ only by the C1/C2 dimorphism defined by positions 77 and 80, which are located on the α1 helix, near the peptide C-terminal amino acid (pΩ) (Fig. 1A, B). We expressed C*08 and C*05 to similar levels in 721.221 cells that are otherwise deficient in classical HLA-I, HLA-A, -B and -C (SFig. 1A). These cells displayed canonical dimorphic recognition by KIR, with KIR2DL2/3 demonstrating strong binding to C*08, and KIR2DL1 demonstrating exclusive binding to C*05 (SFig. 1B). We next used these cells as targets in functional experiments with Jurkat-T cells expressing C*08 restricted KRAS-G12D specific T cell receptor, TCR9a or TCR10. As expected, both the 9mer and 10mer G12D but not their WT KRAS peptides activated their respective TCR9a and TCR10 expressing T cells in the presence of C*08 (Fig. 1C-D) (Sim et al., 2020). Unexpectedly, we observed no activation of TCR9a and only weak activation of TCR10 when the 9mer and 10mer G12D KRAS peptides were presented by C*05, respectively (Fig. 1C-D) (Sim et al., 2020). To confirm this was not due to inefficient peptide loading, we utilized C*08 and C*05 expressing cell lines deficient in transporter associated with antigen presentation (TAP) (SFig. 1C). These experiments yielded similar results using a different functional readout (expression of CD69), as G12D-9mer loaded on C*05 did not activate Jurkat-TCR9a cells, while G12D-10mer loaded on C*05 did stimulate Jurkat-TCR10 cells but required higher peptide concentrations than when loaded on C*08 (SFig. 1C). Thus, the HLA-C1/C2 dimorphism impacts T-cell recognition of KRAS-G12D neoantigen even though both residues 77 and 80 are not in contact with TCR. We next explored if the differences in T-cell recognition of C*08 and C*05 were due to differences in the binding of peptide (Fig. 2) or TCR (Fig. 3) to HLA-C.

**Figure 2.**
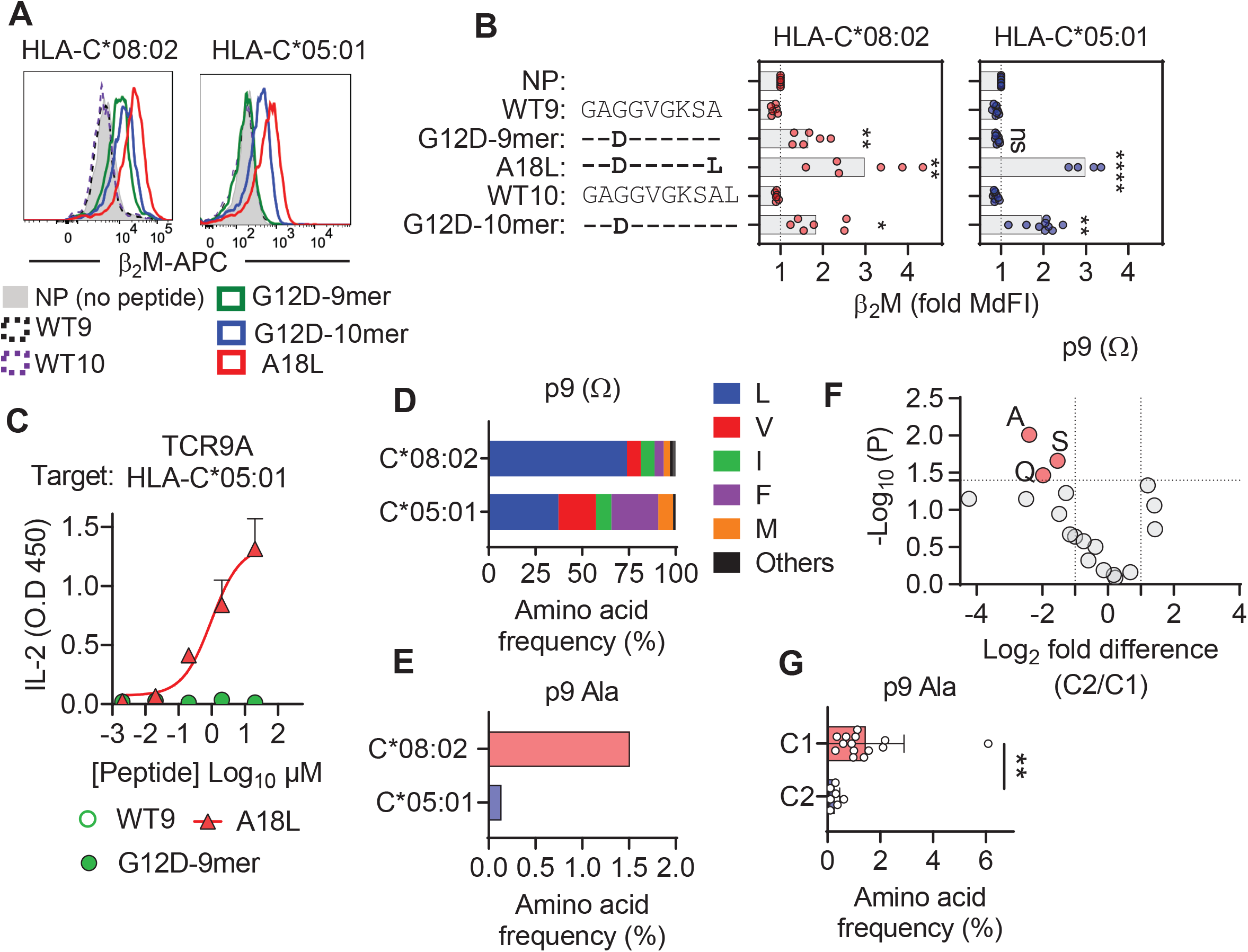
C1 but not C2 HLA-C allotypes present peptides with C-terminal (pΩ) Ala. **(A)** Stabilization of HLA-C on TAP-deficient 221 cells expressing HLA-C*08:02 (left) or HLA-C*05:01 **(B)** Data from A shown as fold median fluorescent intensity (MdFI) relative to no peptide (NP) from a minimum of four independent experiments. **(C)** Stimulation of TCR9a^+^ Jurkat cells by 221-C*05:01 cells preloaded with KRAS peptides at indicated concentrations. Means and standard errors of IL-2 concentration in culture supernatant measured by ELISA from two biological replicates is shown. **(D)** Frequency of indicated residues at the C-terminus (pΩ) in peptides eluted from HLA-C*08:02 or HLA-C*05:01. **(E)** Frequency of pΩ Ala in peptides eluted from HLA-C*08:02 or HLA-C*05:01. **(F)** Volcano plot displaying pΩ amino acid frequency from 21 HLA-C allotypes. The C2/C1 ratio is shown for the average frequency of each amino acid. **(G)** The frequency of pΩ Ala from 21 HLA-C allotypes by C1/C2 status. Statistical significance was assessed by unpaired t-test with Welsh’s correction, * = p<0.05, ** = p<0.001, **** = p<0.0001.

**Figure 3.**
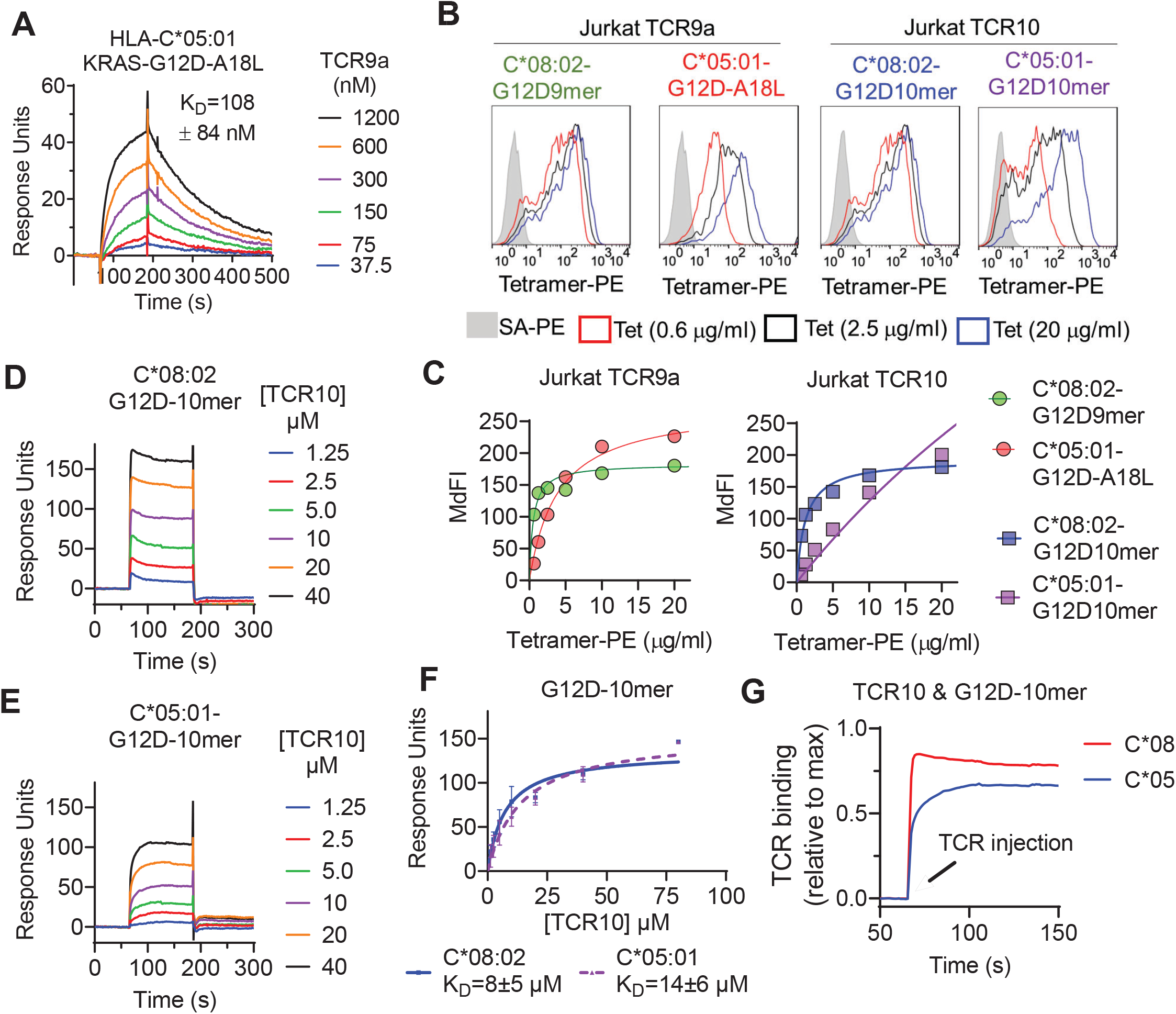
TCR binding is weaker to C2 HLA-C. **(A)** Binding of TCR9a to captured HLA-C*05:01-KRAS-G12D-A18L determined by surface plasmon resonance (SPR). Dissociation constant was determined by kinetic curve fitting of 12 curves from two independent experiments. **(B-C)** Binding of HLA-C*08:02 or HLA-C*05:01 tetramers to Jurkat T-cells expressing TCR9a or TCR10 at indicated tetramer concentrations. HLA-C was refolded with indicated peptides. **(C)** Summary of (B) from two independent experiments. **(D-E)** Binding of TCR10 to captured HLA-C*08:02-G12D-10mer and HLA-C*05:01-G12D-10mer by SPR. **(F)** Non-linear curve fitting of TCR10 binding to G12D-10mer bound to C*08:02 or C*05:01. Dissociation constants were determined by steady state kinetics from two independent experiments. **(G)** Association binding of TCR10 with G12D-10mer presented by C*08:02 or C*05:01. SA = streptavidin, Tet = tetramer.

### KRAS peptide C-terminal residue distinguishes C1 and C2 HLA-C allotypes

To characterize KRAS-G12D peptide binding to C*08 and C*05, we carried out peptide loading assays with TAP-deficient cells expressing C*08 or C*05. Similar to C*08, C*05 was not stabilized by WT-KRAS peptides likely due to the absence of Asp/Glu at p3, a critical anchor for C*08 and C*05 (Fig. 2A, B). However, G12D-9mer only stabilized C*08 and not C*05, while G12D-10mer stabilized both allotypes (Fig. 2A, B). An unusual feature of the G12D-9mer was pΩ Ala, as its short side chain does not fill the pΩ pocket (F pocket), in contrast to G12D-10mer with pΩ Leu (Sim et al., 2020). We previously demonstrated that substitution of pΩ Ala to Leu in G12D-9mer (A18L-9mer: ^10^GA**D**GVGKS**L**) improved binding to HLA-C*08:02, and T-cell recognition by Jurkat-TCR9a (Sim et al., 2020). Similarly, A18L-9mer bound C*05 well (Fig. 2A, B) and stimulated Jurkat-TCR9a cells (Fig. 2C, SFig. 1B). Thus, the inability of G12D-9mer to bind C*05 accounts for its lack of activation of Jurkat-TCR9a cells. Further, these data suggest that the C1/C2 dimorphism selectively impacts peptide binding to HLA-C in a sequence dependent manner.

To explore if pΩ Ala is a general feature of peptides that bind C*08 and not C*05, we examined their immunopeptidomes using two publicly available datasets (see methods) (Di Marco et al., 2017; Sarkizova et al., 2020). The residues that form the pΩ binding pocket of C*08 and C*05 are identical and accordingly the five most abundant pΩ residues (Leu, Val, Ile, Phe and Met) were the same in both allotypes, accounting for 97% of C*08 and 98% of C*05 sequences (Fig. 2D). Leucine was the most common pΩ in both allotypes but had a higher frequency in C*08 peptides than C*05 (C*08=73%, C*05=37%) (Fig. 2D). The significance of this difference is unknown, however peptides with pΩ Leu, Val, Ile, Phe and Met bind well to both C*08 and C*05 (Sim et al., 2017). Among 9mer peptides, the frequency of Ala at pΩ position was 1.5% (30 out of 1986) in C*08, significantly higher (p = 0.0016 Fishers exact test) than that of 0.13% (1 out of 732) in C*05 bound peptides (Fig. 2E). Similar results were observed for 10mers, where pΩ Ala was more common among C*08 peptides (5/349, 1.4%) than C*05 peptides (0/152, 0%) (SFig. 2). We next analyzed the pΩ amino acid frequency amongst 9mer sequences eluted from 14 C1 allotypes and 7 C2 allotypes. In concordance with the comparison of C*05 and C*08 immunopeptidomes, pΩ Ala was the most enriched in C1 allotypes, with a frequency of 1.45±1.4% in C1 allotypes compared to 0.27±0.19% in C2 allotypes (Fig. 2F, G). In addition, Gln and Ser were significantly enriched at pΩ of C1 allotypes, but at lower frequencies than Ala. Thus, C1 allotypes bind peptides with pΩ Ala to a greater extent than C2 allotypes, consistent with the ability of G12D-9mer to bind C*08 and not C*05.

### Impact of C1/C2 dimorphism on TCR binding to HLA-C

We next explored the impact of the C1/C2 dimorphism on TCR binding to HLA-C using recombinant HLA-C refolded with individual peptides. The solution binding affinities of the two neoantigen specific TCRs, TCR9a and TCR10, to their cognate peptides presented by C*08 or C*05 were determined by surface plasmon resonance (SPR) using recombinant TCRs as analytes. As G12D-9mer with pΩ Ala does not bind C*05 (Fig 2A), we chose the Leu anchor variant A18L-9mer as the peptide for C*05 to ensure HLA-C stability. While TCR9a bound to the C1 allotype HLA-C*08 in the presence of G12D-9mer peptide with 16nM affinity (Sim et al., 2020), it displayed a binding affinity of 108 nM for the C2 allotype HLA-C*05 in the presence of Leu anchor variant of G12D-9mer, a ∼7-fold affinity reduction compared to the C1 allotype (Fig. 3A). Consistent with their solution binding affinities, the C*08-G12D-9mer tetramers bound TCR9a expressing Jurkat cells with lower EC_50_ than C*05-A18L-9mer tetramers (Fig 3B,C). This reduced binding affinity and avidity may explain the reduced sensitivity of Jurkat-TCR9a cells for A18L-9mer loaded on C*05 compared to C*08 (SFig 3). In solution, TCR10 bound both the C1 and C2 allotypes with μm affinities (Fig 3D-F), though slightly weaker affinity was observed with C*05. By steady-state kinetics, TCR10 binding was less than two-fold weaker to C*05:01-G12D-10mer (K_D_ = 13.5±6 μM) than C*08:02-G12D-10mer (K_D_ =8±5 μM) (Fig. 3D,E). However, by kinetic analysis of the binding curves (Fig. 3G), TCR10 had almost twenty-fold slower on rate with C*05:01-G12D-10mer (K_on_ = 6.2±1.2×10^3^ M^-1^s^-1^) than with C*08:02-G12D-10mer (K_on_ = 1.2± 0.9×10^5^ M^-1^s^-1^). Consistently, C*08-G12D-10mer exhibited better binding to TCR10-expressing Jurkat T cells than C*05-G12D-10mer, which failed to reach binding saturation at the highest concentration of tetramer tested (40 μg/ml) (Fig 3B,C). Thus, both TCR9a and TCR10 exhibited better binding to the C1 allotype than C2.

Neither TCR9a nor TCR10 make direct contacts to HLA-C positions Ser 77 and Asn 80 (Sim et al., 2020). To understand the impact of C2 residues Asn 77 and Lys 80 on T cell recognition, we determined the crystal structure of TCR9a in complex with C*05-A18L-9mer to a high resolution of 1.9 Å (Supplementary Table 1). TCR9a adopted the same binding mode when in complex with C*05, as in the C*08 complex structure (Fig. 4A). TCR9a recognition of C*05-A18L-9mer was via CDR3α Q98 that forms hydrogen (h) bonds with Arg 156 and Gln 157 of HLA-C and CDR2β Tyr48 and Glu49 that coordinate peptide p7 Lys (Fig. 4B, C). CDR3β of TCR9a E95 formed a salt bridge with C*05 Arg 69 (Fig. 4D). All these contacts observed between TCR9a and the C2 HLA-C*05 are conserved in the structure of TCR9a complexed with C*08-(Supplemental Table 2). Like the C1 HLA-C*08, the dimorphic residues 77 and 80 on the C2 HLA-C*05 also do not interact with TCR9a. The closest TCR residue to HLA-C position 77 or 80 is Glu 49 of the TCRβ chain which interacts with Lys p8 of the peptide but is greater than 7Å away from HLA-C position 80. Docking TCR10 onto a structure of C*05:01-G12D-10mer using our TCR10-C*08:02-G12D-10mer complex, revealed no obvious impact on TCR contacts (SFig. 4) (Bai et al., 2021; Sim et al., 2020), suggesting the dimorphic positions on C*05 do not directly contact TCR.

**Figure 4.**
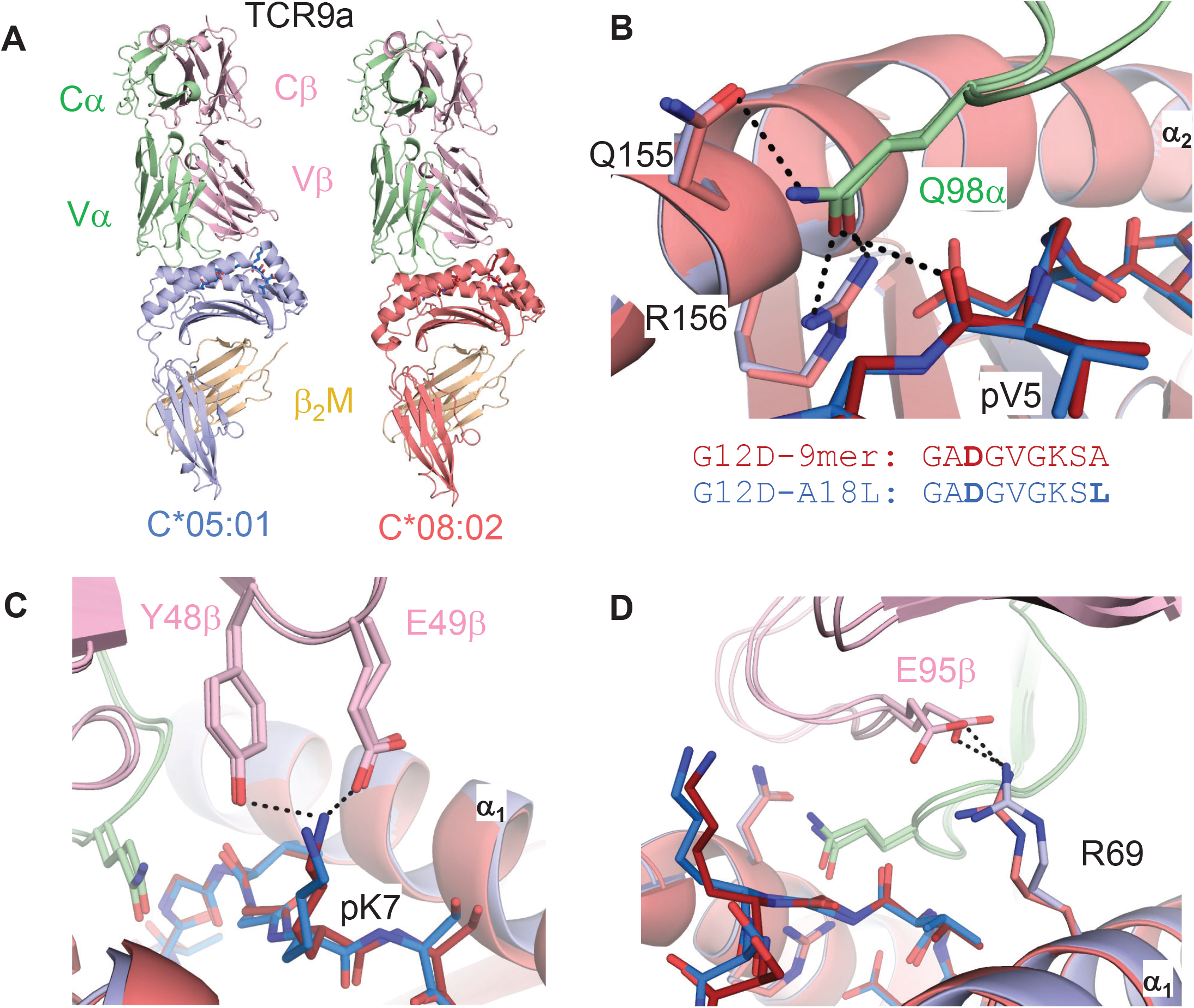
Minimal impact of C1/C2 dimorphism on TCR:HLA-C complex structure. **(A)** Side view of TCR9a in complex with HLA-C*05:01-G12D-A18L (*left*) and HLA-C*08:02-G12D-9mer (*right;* PDB:). **(B-D)** Interactions of TCR9a with HLA-C*05:01-A18L (blue) and HLA-C*08:02-G12D-9mer (red). TCR9a α chain, green, β chain, pink; β2-microglobulin (β_2_M), gold.

### The C1/C2 dimorphism affects pΩ-1 backbone position and sidechain preference

The lack of direct contacts between TCR and the HLA-C1/C2 dimorphic residues raised the question if the C1/C2 dimorphism indirectly affected TCR binding through peptide conformation. The position 77 side chain is orientated directly into the PBG, while position 80 sits atop the α1-helix (Fig. 1B). Position 77 is Ser in C*08 (C1) and Asn in C*05 (C2) and both form a h-bond of the same distance with the amide of the terminal peptide bind between pΩ and p8 (pΩ-1), (Fig. 5A). Comparing the structures of C*05 and C*08, the HLA-C α_1_-helices were displaced relative to each other and peptide p8 was higher out of the PBG in C*05 compared to C*08 (Fig. 5B-D). This displacement was measured by comparing the distance between the Cα of peptide p8 (Ser) with that of the Cβ of HLA-C position 77 (the first carbon of the position 77 side chain). This distance was 4.7Å and 5.7Å in the C*08 and C*05 structures, respectively (Fig. 5B&C). We next examined this distance in ten additional published HLA-C structures, four C1 (HLA-C*03:04, PDB:1EFX, HLA-C*07:02, PDB:5VGE, HLA-C*08:01, PDB:4NT6, HLA-C*08:02-G12D-10mer, PDB: 6ULK) and six C2 (HLA-C*04:01, PDB:1QQD, HLA-C*05:01-G12D-10mer, PDB: 6JTO, HLA-C*05:01, PDB: 5VGD, HLA-C*06:02, PDB:5W6A, 5W69, 5W67). We observed that the difference between the Cα of peptide p8 and HLA-C 77 Cβ was maintained between C1 and C2 HLA-C structures (C1=4.6±0.11Å, C2=5.7±0.22Å) (Fig. 5G). In contrast, a conserved contact with the peptide N terminus and Tyr171 Oγ was the same between C1 and C2 HLA-C structures (2.70 Å vs 2.76 Å) (Fig. 5E&H). A consequence of this displacement is the proximity of the p8 side chain to the HLA-C α_1_ helix (Fig. 5F). We compared the distance between the p8 side chain and HLA-C Val 76 in the 10 HLA-C structures. In all five C1 allotype structures, the p8 side chain was within van der waals range of Val 76 (3.3-4.0Å). In contrast, the p8 side chain was out of contact range with Val 76 in all but one C2 HLA-C structures (Fig. 5F&I). This appears to be due in part to the torsion angle of the terminal peptide bond that orientates p8 towards the α_1_ helix in C1 allotypes and out of the groove in C2 allotypes (Fig. 5J). Thus, the C1/C2 dimorphism impacts the position and orientation of the peptide at pΩ-1, suggesting position 77 is the major determinant of the difference between C*05 and C*08. To interrogate the role of positions 77 and 80 individually, single substitutions in C*05 were made to that of C*08. Cells expressing C*05, C*05 Asn77Ser (N77S), C*05 Lys80Asn (K80N) and C*08 loaded with G12D-9mer or G12D-10mer were used as targets for Jurkat-TCR9a and Jurkat-TCR10 cells respectively. The lack of stimulation of Jurkat-TCR9a by C*05, was minimally impacted by C*05-K80N, while C*05-N77S recovered Jurkat-TCR9a responses, though not completely (Fig. 5K). In contrast, stimulation of Jurkat-TCR10 cells was not impacted by single substitutions at positions 77 and 80, suggesting a synergistic effect of both positions 77 and 80 on recognition of HLA-C by TCR10.

**Figure 5.**
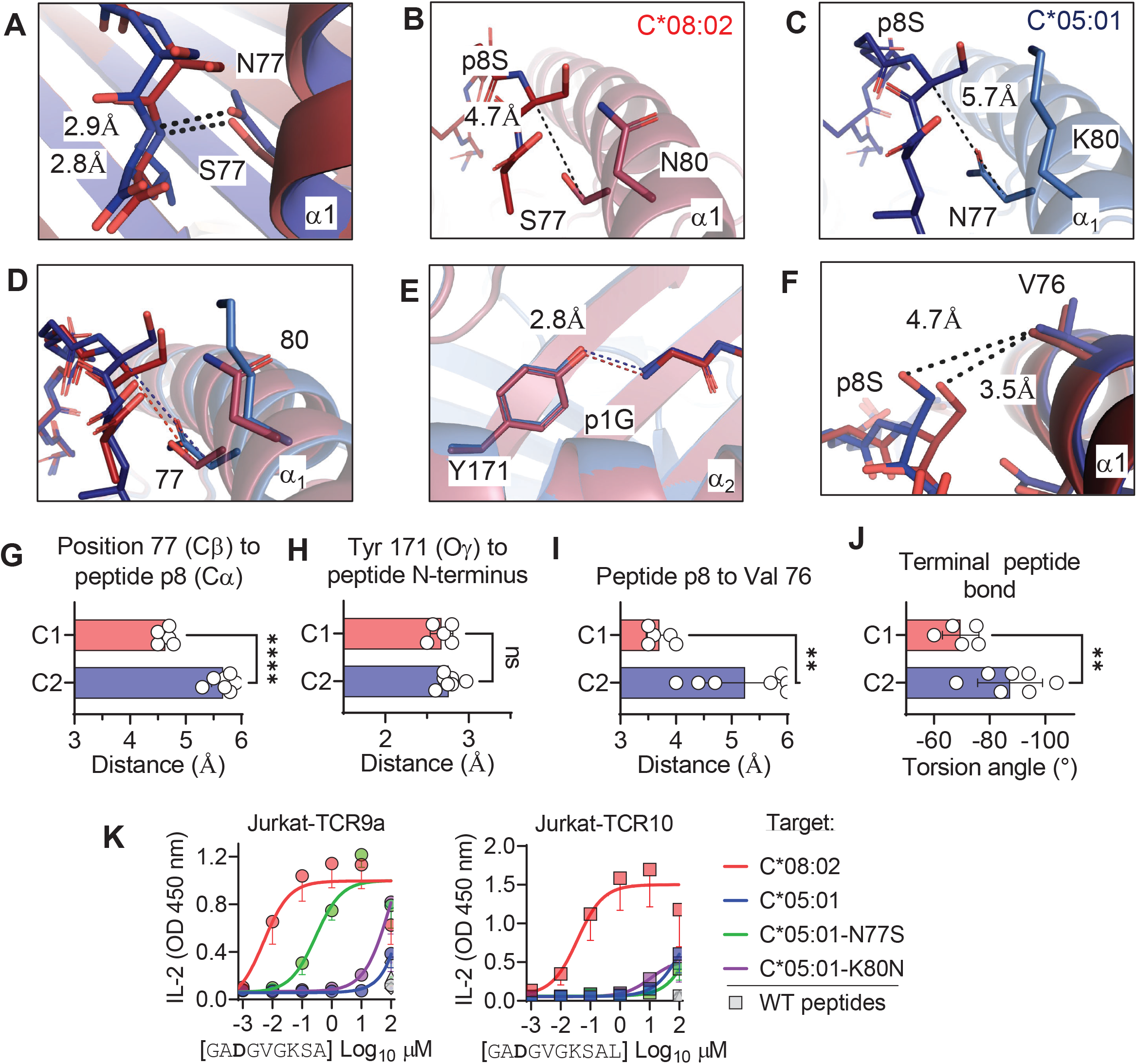
C1/C2 dimorphism determines the distance between peptide p8 and HLA-C. Hydrogen bond between HLA-C position 77 and amide in terminal peptide bond in C*08:02-G12D-9mer and C*05:01-A18L-9mer structures. **(B-D)** Distance between HLA-C position 77 Cβ and Cα of peptide p8 Ser of HLA-C*08:02-G12D-9mer **(B)**, HLA-C*05:01-G12D-A18L **(C)** and both overlaid **(D). (E)** Distance between Tyr 171 Oγ and peptide N-terminus. **(F)** Distance between peptide p8 side chain and HLA-C position 77 Val side chain. **(G)** Distances between HLA-C position 77 Cβ and Cα of peptide pΩ-1 in 12 HLA-C crystal structures. **(H)** Distance between Tyr 171 Oγ and peptide N-terminus in 12 HLA-C crystal structures. **(I)** Distance between peptide pW-1 side chain and HLA-C position 77 Val side chain in 10 HLA-C structures. **(J)** Torsion angle of terminal peptide bond in 12 HLA-C structures **(K)** Stimulation of Jurkat-TCR9a^+^ and Jurkat-TCR10^+^ by 221 cells expressing HLA-C*08:02, HLA-C*05:01, HLA-C*05:01-N77S, or HLA-C*05:01-K80N preloaded with G12D-9mer (*left*) or G12D-10mer (*right*) Means and standard errors of IL-2 concentration in culture supernatant measured by ELISA from three biological replicates are shown. Significance in G-J was measured using an unpaired t test with Welch’s correction, ** = p<0.01, **** = p<0.0001.

We next explored if the shift in pΩ-1 position is associated with changes in the HLA-C immunopeptidome other than those at pΩ (Fig. 2). Analyzing 9mer sequences unique to C*08 and C*05, we compared the amino acid frequency at each position (1-9) by Pearson correlation. Strong correlations (r > 0.9) were observed at positions 1-3, while the pΩ was slightly weaker (r = 0.8), reflecting the small differences in pΩ usage (Fig. 6A, SFig. 5A). In contrast, there was no correlation in amino acid usage (r = -0.01) at p8 (pΩ-1)(Fig. 6A). At pΩ-1, Ala and Ser made up over 30% of sequences unique to C*08, but only 5% of sequences unique to C*05. In contrast, Leu, Lys and Arg were found in approximately 40% of sequences unique to C*05, but only 7% of sequences unique to C*08 (Fig. 6A). We observed similar findings for p9 (pΩ-1) in an analysis of 10mers (SFig. 5B). Next, we analyzed the relative amino acid frequencies at p8 of 9mers eluted from 14 C1 allotypes and 7 C2 allotypes (Fig. 6B). We found that at p8, C1 allotypes are enriched for Ser, Ala and Pro, while C2 allotypes are enriched for Phe, Leu, Lys, Met, Arg, Trp and Tyr (Fig. 6B, SFig. 5C). There was a strong correlation between amino acid enrichment at p8 in C2 allotypes and amino acid side chain volume, demonstrating that the C1/C2 dimorphism imposes a size constraint on the peptides bound to HLA-C (Fig. 6C). In peptides eluted from C1 allotypes, the preference for smaller residues at p8 was accentuated in peptides with pΩ-Ala compared to those with pΩ-Leu (SFig.5D), suggesting that size at pΩ-1 is especially important for peptides with pΩ-Ala.

**Figure 6.**
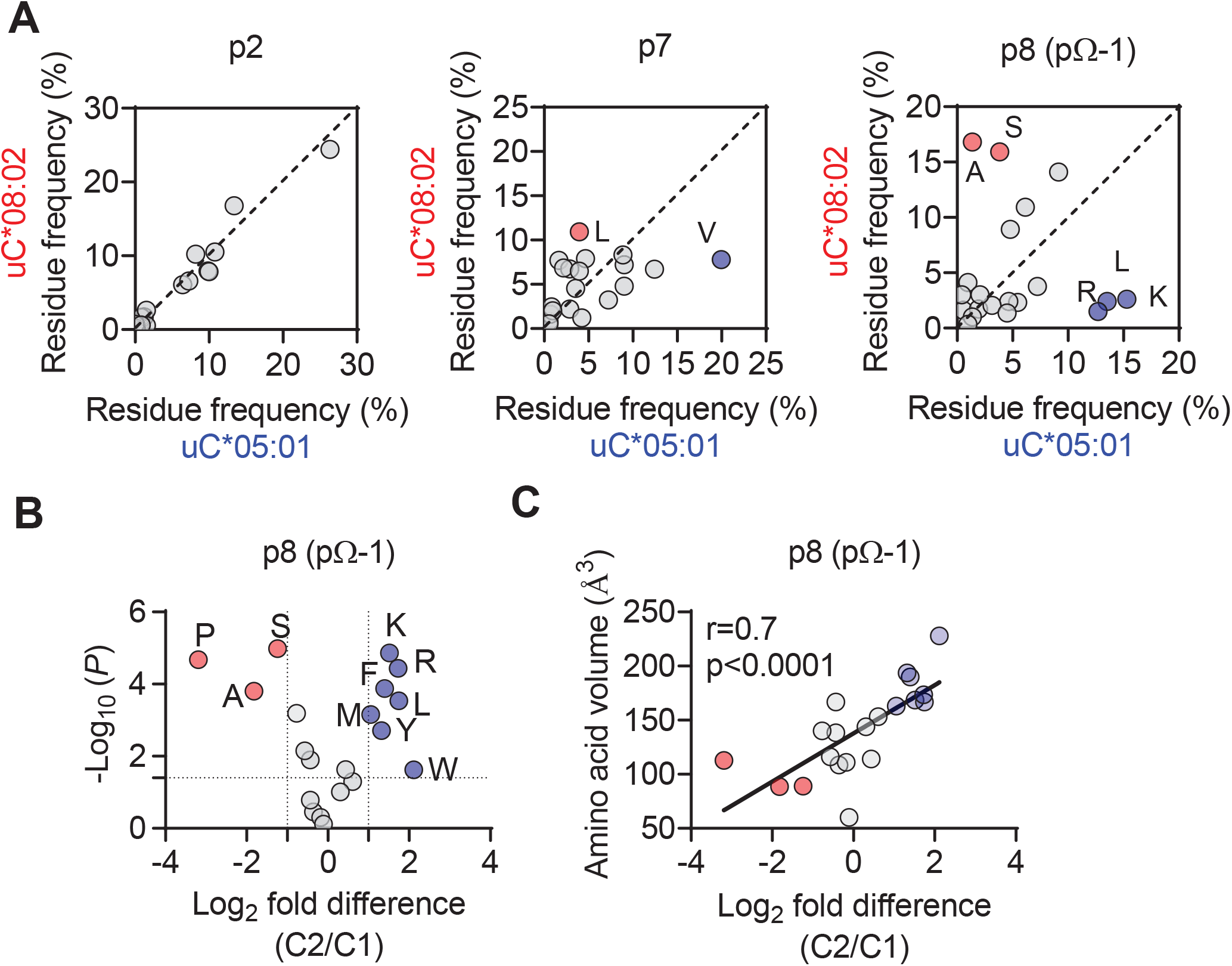
The C1/C2 dimorphism selects for side chain size at pΩ-1. **(A)** Correlation of amino acid frequency at p2, p7 and p8 (pΩ-1) of 9mer peptides unique to HLA-C*08:02 (uC*08:02) and unique to HLA-C*05:01 (uC*05:01). **(B)** Volcano plot displaying enrichment at p8 amino acid frequency from peptides eluted from 14 HLA-C allotypes based on C1/C2 status. Amino acids with two-fold enrichment and statistically significant differences (p<0.05) determined by student t-test are shown. A total of 26,543 peptide sequences were included. **(C)** Correlation of amino acid volume with amino acid enrichment at p8 of peptides eluted from HLA-C defined by C2/C1 status.

### Large amino acid side chains at peptide pΩ-1 position impair TCR9a recognition

Finally, we tested the importance of side chain size at pΩ-1 by substituting p8 Ser in G12D-9mer and A18L-9mer to Ala, Gly, Glu, Val, Leu, Lys and Arg. Stimulation of Jurkat-TCR9a by C*08 cells loaded with G12D-9mer p8 substitutions to Leu, Lys and Arg substantially increased the EC_50_ by approximately 1000-fold, while substitution to Ala, Gly, Glu, Val had a minimal impact on the EC_50_ (Fig. 7A-C). The same p8 substitutions in A18L-9mer had a much less dramatic impact on Jurkat-TCR9a stimulation, however p8 Lys, Arg and Leu increased the EC_50_ by roughly 10-fold (Fig 7D-F). There was not a strict correlation between Jurkat-TCR9a stimulation and stabilization of HLA-C on TAP-deficient C*08^+^ cells (SFig. 6). For example, in G12D-9mer, S8L was a poor ligand for Jurkat-TCR9a, but stabilized HLA-C better than S8G which was a better ligand for Jurkat-TCR9a. Further, substitution at p8 in A18L-9mer had a minimal effect on HLA-C stabilization, but still impacted Jurkat-TCR9a activation (Fig. 7F, SFig. 6). This may be due in part to differences in sensitivity between these assays but also suggests a complex relationship between p8 sequence, HLA-C stabilization and T-cell recognition. Inline with our other data, the impact of p8 size demonstrates how molecular changes outside TCR contacts can directly impact T cell recognition.

**Figure 7.**
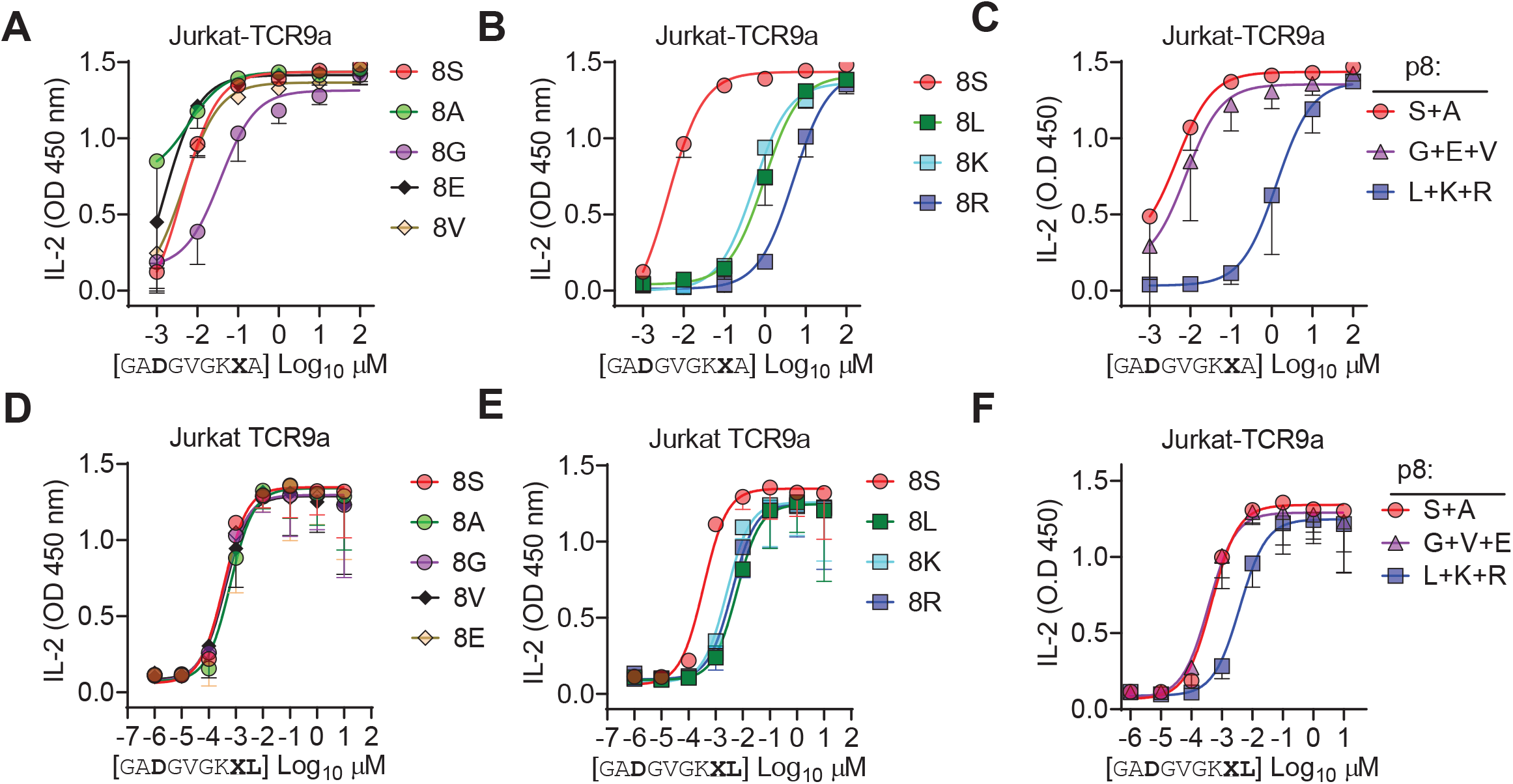
Large residues at p8 diminish T-cell recognition of C1 HLA-C. **(A-B)** Stimulation of Jurkat-TCR9a^+^ cells by 221-C*08:02 cells preloaded with pΩ-1 substitutions of G12D-9mer (**A-C**) and A18L-9mer (**D-F**) at indicated concentrations. Each peptide was tested individually in two independent experiments and displayed by p8 sequence as indicated. Means and standard errors of IL-2 concentration in culture supernatant measured by ELISA are shown. Data from A&B are summarized in C, data from D&E are summarized in F.

## Discussion

HLA-I molecules form ligands for many immunoreceptors and interpreting disease associations with HLA-I requires modeling interactions with different immune populations (Debebe et al., 2020). The C1/C2 dimorphism in HLA-C plays a critical role in innate immunity by forming ligands for the NK cell receptors KIR2DL1 and KIR2DL2/3. By comparing T cell recognition of two HLA-C allotypes identical in sequence except for the C1/C2 dimorphism, we assessed how this dimorphism affects adaptive immunity. We examined two clinically relevant, HLA-C*08:02 restricted neoantigens, G12D-9mer and G12D-10mer derived from the oncogenic hotspot mutation KRAS-G12D. We reject the null hypothesis that the C1/C2 dimorphism exhibits no effect on T cell recognition as C*05 was a poor ligand for TCR9a^+^ and TCR10^+^ T cells. Our study identified a clear role for the C1/C2 dimorphism on peptide binding to HLA-C; specifically as G12D-9mer only bound C*08 and not C*05, but also in more general terms via analysis of HLA-C immunopeptidomes. Further, we identified an impact of the C1/C2 dimorphism on TCR binding as TCR9a and TCR10 binding to C*05 was weaker than binding to C*08.

By investigating the exclusive restriction of TCR9a to C*08, we uncovered a novel role for the C1/C2 dimorphism in peptide presentation. Specifically, the C1/C2 dimorphism defines two distinct PBG structures that impose size constraints at pΩ-1 and allow C1 but not C2 allotypes to bind peptides with shorter pΩ anchors, like Ala in G12D-9mer. The primary cause of these distinct PBGs is likely the size of position 77, which forms a conserved h-bond with the amide of pΩ in both C1 and C2 HLA-C. As Asn (C2) is larger than Ser (C1), it results in peptide pΩ-1 protruding out of the groove thus permitting larger amino acids at pΩ-1 in C2 allotypes compared to C1. In C1, pΩ-1 sits lower in the groove, restricting pΩ-1 side chains to smaller residues to avoid clashing with HLA-C position Val 76. Consistently, substitutions at p8 in G12D-9mer to the larger Leu, Lys or Arg substantially reduced C1 recognition by TCR9a, while substitution to Gly had only a minor decrease. Further, the penalty imposed for large residues at p8 was diminished with A18L-9mer, suggesting a canonical pΩ anchor can accommodate unfavorable p8 residues, likely via orientating p8 side chain out of the groove like peptides in C2 allotypes. In C1 allotypes, the additional contacts between Val 76 and p8 may explain the ability of C*08:02 to bind peptides with short, suboptimal pΩ anchors. In contrast, pΩ-1 in C2, which is higher in the PBG and orientated away from the α1 helix, is unable to contact Val 76 and therefore cannot support peptides with pΩ Ala. Consistently, substitution of Asn 77 to Ser in C*05 substantially recovered TCR9a recognition of G12D-9mer.

In addition to peptide binding to HLA-C, TCR binding affinity and T cell sensitivity was reduced in the presence of C2 residues. Neither N77S or K80N substitutions improved the sensitivity of TCR10 for C*05 and G12D-10mer, suggesting a synergistic effect of the C2 residues on reducing TCR recognition. TCR10 displayed a much slower on-rate in binding C*05-G12D-10mer and even after controlling for a poor pΩ anchor, TCR9a was more sensitive to A18L-9mer with C*08 than C*05. Comparing our TCR9a-C*05:01-A18L-9mer structure with that of TCR9a:C*08:02-G12D-9mer revealed no impact on TCR contacts. Further, modelling TCR10 in complex with a recently solved crystal structure of C*05-G12D-10mer revealed no impact on TCR contacts (6JTO) (Bai et al., 2021). The most likely explanation for diminished TCR binding in the presence of C2 residues is that of conformational flexibility. Clearly, TCR9a and TCR10 can bind C*05, however the ability to make the same contacts with C*08 in C*05 likely requires more energetically unfavorable conformational changes, in either peptide, HLA-C or both, exemplified by the slow on-rate of TCR10 with C*05-G12D-10mer. Indeed, recent evidence suggests that TCR recognition can be optimized by peptide substitutions that allow TCR contacts to form more readily (Devlin et al., 2020; Duru et al., 2020). It is likely that the converse is also true, slight alterations in amino acid side chain orientations or flexibility, may impose a higher energy barrier for efficient TCR recognition.

The G12D-9mer and G12D-10mer are considered public or shared neoantigens due to the high frequency of cancers with KRAS-G12D mutations (Cox et al., 2014; Pearlman et al., 2021; Stephen et al., 2014). Therefore, the small differences between C*08 and C*05 investigated here have clinical implications for mounting an effective anti-tumor immune response (Tran et al., 2016). The ability of C*05 to bind G12D-10mer means HLA-C*05:01 positive individuals may be eligible for such therapies, however identifying a more sensitive receptor than TCR10 would be needed to translate these findings for clinical application.

The consequences of our findings may have broad implications for T cell recognition of HLA-C beyond the specific example of KRAS-G12D recognition in cancer. For example, in infectious diseases, viral escape mutants may eliminate T cell recognition of C1 allotypes by increasing side chain size at pΩ-1. Similarly, escape mutants that substitute pΩ to smaller residues like Ala may escape T cell recognition of C2 HLA-C but not C1. For peptide vaccines, pΩ-1 size could be optimized depending on whether the peptide targets a C1 or C2 HLA-C allotype. A reasonable prediction from our work is that if a T cell epitope contains a canonical pΩ, it will be immunogenic in individuals carrying either C*08 and C*05. Indeed, there is evidence that HIV infected individuals can respond to the same epitope (SAEPVPLQL) if they carry HLA-Cw5 or HLA-Cw8 (Addo et al., 2001). However, it is possible that preference for pΩ-1 size could engender differential T cell immunity based on differences in HLA-C and TCR binding. These effects need to be investigated in further experiments of HLA-C restricted T cells. It is worth noting that the dimorphic HLA-C response to TCR is intrinsic to the structure of HLA-C and thus may also impact interactions with other immunoreceptors. In particular, the differences in amino acid preferences at pΩ-1 may have enormous consequences for KIR recognition of HLA-C, as this residue is central to the KIR binding site (Boyington et al., 2000; Boyington and Sun, 2002; Fan et al., 2001). Our study highlights how small HLA polymorphisms can impact interactions with both peptide and immunoreceptors with implications for immune responses and cancer immunotherapy. Finally, our study establishes that T cells, like NK cells, can discriminate between the dimorphic HLA-C allotypes.

## Materials and methods

### Cell lines and culture

Cell lines were cultured in IMDM and 10% fetal calf serum (FCS) at 37°C and 5% CO_2_. 721.221 (221) cells expressing HLA-C*05:01 and the TAP inhibitor ICP47 (221-C*05:01-ICP47) were described previously (Sim et al., 2017; Sim et al., 2019). Cas9 expressing 221 cells were a generated by lentiviral transduction. cDNA encoding full-length HLA-C*08:02 was expressed in 221-Cas9 cells by retroviral transduction with vectors on the pbabe backbone. Two TAP1 gRNAs (Genscript, USA) were introduced by lentiviral transduction. Jurkat T cells expressing TCR9a and TCR10 were previously described (Sim et al., 2020).

### Peptide loading assay

Peptide loading assays were performed as described (Sim et al., 2017; Sim et al., 2019). 221-C*05:01-ICP47 and 221-Cas9-C*08:012-TAP cells were cultured overnight at 26°C with 100 μM of peptide. Cells were then stained with anti-HLA-I mAb (APC, W6/32, Biolegend USA, #311410). Expression of HLA-C was determined by flow cytometry. Each peptide was tested at least twice in independent experiments. Peptides were synthesized at >95% purity (Genscript, USA).

### T cell activation assay

Peptides were incubated with 10^5^ 221-HLA-C^+^ target cells for 4hrs at 37°C before incubation with 10^5^ Jurkat-TCR^+^ cells overnight. The following day, cell culture supernatants were recovered and IL-2 measured by ELISA using the ELISA MAX™ Deluxe Set Human IL-2, Biolegend, USA (#431804). For assays with TAP-deficient target cells, 10^5^ target cells were incubated overnight at 26°C with peptide ranging from 1 μM to 0.01 nM. The following day, target cells were mixed with TCR^+^ Jurkat T cells for 6 hr at 37°C. Cells were then washed twice in PBS and stained with mAbs to CD69 (APC, FN50, BD Biosciences USA, #555533) and CD3 (APC-Cy7, UCHT1,

Biolegend USA, #300426). Expression of CD69 was measure on CD3^+^ cells by flow cytometry. All peptides were tested at indicated concentrations at least twice in independent experiments.

### HLA-C immunopeptidomics

For comparing the immunopeptidomes of HLA-C*05:01 and HLA-C*08:02, 8mer, 9mer and 10mer sequences from two studies were combined and duplicates removed (Di Marco et al., 2017; Sarkizova et al., 2020). A total of 1928 C*05 and 3182 C*08 non-redundant 9mer sequences were compared of which 1196 sequences were shared between C*05 and C*08 (SFig. 5). To compare C*05 and C*08 immunopeptidomes, the analysis included 732 and 1986 9mer sequences unique to C*05 and C*08, respectively. For comparing the immunopeptidomes of C1 and C2 HLA-C allotypes, 9mer peptides eluted from 21 different HLA-C allotypes were obtained from two published studies (Di Marco et al., 2017; Sarkizova et al., 2020). The C1 HLA-C allotypes were; HLA-C*01:02, C*03:02, C*03:03, C*03:04, C*07:01, C*07:02, C*07:04, C*08:01, C*08:02, C*12:02, C*12:03, C*14:02, C*14:03 and C*16:01. The C2 HLA-C allotypes were; HLA-C*02:02, C*04:01, C*04:03, C*05:01, C*06:02, C*15:02 and C*17:01. For C*03:02, C*04:03, C*07:04 C*08:01, C*12:02, C*14:03, data were only available from one study (Sarkizova et al., 2020). For the remaining allotypes, sequences from both studies were combined and duplicates removed. A total of 18275 peptides from C1 allotypes and 8268 peptides from C2 allotypes were studied. The amino acid frequency at positions 8 (pΩ-1) and 9 (pΩ) were determined for each allotype independently and collected to derive an average amino acid frequency for C1 and C2 allotypes. The relative amino acid frequency between C2 and C1 allotypes (C2/C1 fold difference) was determined for each amino acid and significant differences determined by two-sided student t-test.

### Protein expression, purification and crystallization

HLA-C*05:01 and TCR proteins were produced largely as described (Clements et al., 2002; Garboczi et al., 1992; Sim et al., 2020; Tikhonova et al., 2012). DNA encoding residues 1-278 of HLA-C*05:01, 1-99 of β_2_M and TCR alpha (1-208) and beta (1-244) chains were synthesized and cloned into the bacterial expression vector pET30a via NdeI and XhoI (Genscript, USA). Proteins were expressed as inclusion bodies in BL21 (DE3) cells (Invitrogen, USA) and dissolved in 8M urea, 0.1 M Tris pH 8. Proteins were refolded by rapid dilution in 0.5 M L-Arginine, 0.1M Tris pH 8, 2 mM EDTA, 0.5 mM oxidized Glutathione, 5 mM reduced Glutathione. Proteins were purified by ion-exchange chromatography (Q column, GE Healthcare, USA) followed by size exclusion chromatography with a Superdex 200 column (GE Healthcare, USA). HLA-C*05:01-G12D-A18L-9mer and TCR9a were concentrated to 10 mg/ml and crystals grown under the same conditions as TCR9a/d with HLA-C*08:02-G12D-9mer (Sim et al., 2020) and were 22% PEG 3350, 0.1 M MOPS pH 7.1 and 0.25 M MgSO_4_.

### Data collection, structure determination and refinement

Crystals were immersed in cryoprotectant (crystallization condition plus 20% glycerol) before flash-cooling in liquid nitrogen. A single data set was collected on the SER-CAT 22 ID beamline (Argonne National Laboratory, IL, USA), processed and merged using HKL2000 (Otwinowski and Minor, 1997). TCR9a:HLA-C*05:01-G12D-A18L-9mer structure was solved by molecular replacement method with Phaser in the CCP4 package using the TCR9a:HLA-C*08:02-G12D-9mer complex as a search model (PDB: 6ULN), (Sim et al., 2020). The model was built and refined with Coot and Phenix (Adams et al., 2002; Collaborative Computational Project, 1994; Emsley and Cowtan, 2004; McCoy et al., 2007). The complex contained one complex per asymmetric unit and belonged to the P2_1_ space group. Peptide and CDR loops were added manually using 2Fo-Fc electron density maps. Graphical figures were generated in PyMOL.

### Surface plasmon resonance (SPR)

SPR was performed largely as described (Sim et al., 2020) with a BIAcore 3000 instrument and analyzed with BIAevalution software v4.1 (GE Healthcare, USA). The pan HLA-I specific mAb W6/32 (Biolegend, USA) was immobilized to CM5 chips (GE Healthcare, USA) at 5000-7000 response units (RU) by primary amine-coupling with a 2 μl/min flow rate in 10mM sodium acetate pH 5.5. HLA-C was captured by W6/32 at 400 - 700 RU in PBS. Soluble TCR heterodimers were used as analytes in 10 mM HEPES pH 7.5 and 0.15 M NaCl with a flow rate of 50 μl/min. TCRs were injected for two minutes followed by a dissociation of ten minutes. Binding was measured with serial dilutions of TCR from 80 μM to 1.25 μM for TCR10 and 1200 nM to 37.5 nM for TCR9a. Dissociation constants were obtained by modelling steady state kinetics and kinetic curve fitting for TCR9a with BIAevaluation software.

### Flow cytometry

Flow cytometry was performed on an LSR II or Fortessa X20 (BD Biosciences, USA) and a Cytoflex S (Beckman Coulter, USA). Data were analyzed using FlowJo software (Treestar V10, USA). Cytometer setup and tracking beads were run daily and single mAb stained beads were used to determine compensation settings for multicolor experiments

### Statistical analysis

Statistical analyses were carried out in GraphPad PRISM (version 8) and Microsoft Excel.

### Protein Database deposition and files

The C*05:01-A18L-9mer-TCR9a complex was assigned the PDB code: XXXX. The following PDB were used in this paper; 6ULI, 1EFX, 5VGE, 4NT6, PDB: 6ULK, 1QQD, 6JTO, 5VGD, 5W6A, 5W69, 5W67.

## Acknowledgements

We thank Dr. Sumati Rajagopalan for critical reading of the manuscript. We thank Dr. Ludmila Krymskaya of the LIG Flow Core for assistance with cell sorting and staff at Argonne National Laboratory for assistance with collecting x-ray diffraction data. This work was supported by the Intramural Research Program of the NIH, National Institute of Allergy and Infectious Diseases.

## Author contributions

M.J.W.S. and P.D.S. designed research; M.J.W.S., Z.S., J.L., and P.B. performed research; J.L., E.O.L. contributed new reagents/analytic tools; M.J.W.S. and J.L. analyzed data; and M.J.W.S., E.O.L., and P.D.S. wrote the paper. All authors read and commented on the final manuscript.

---

**SFigure 1.**
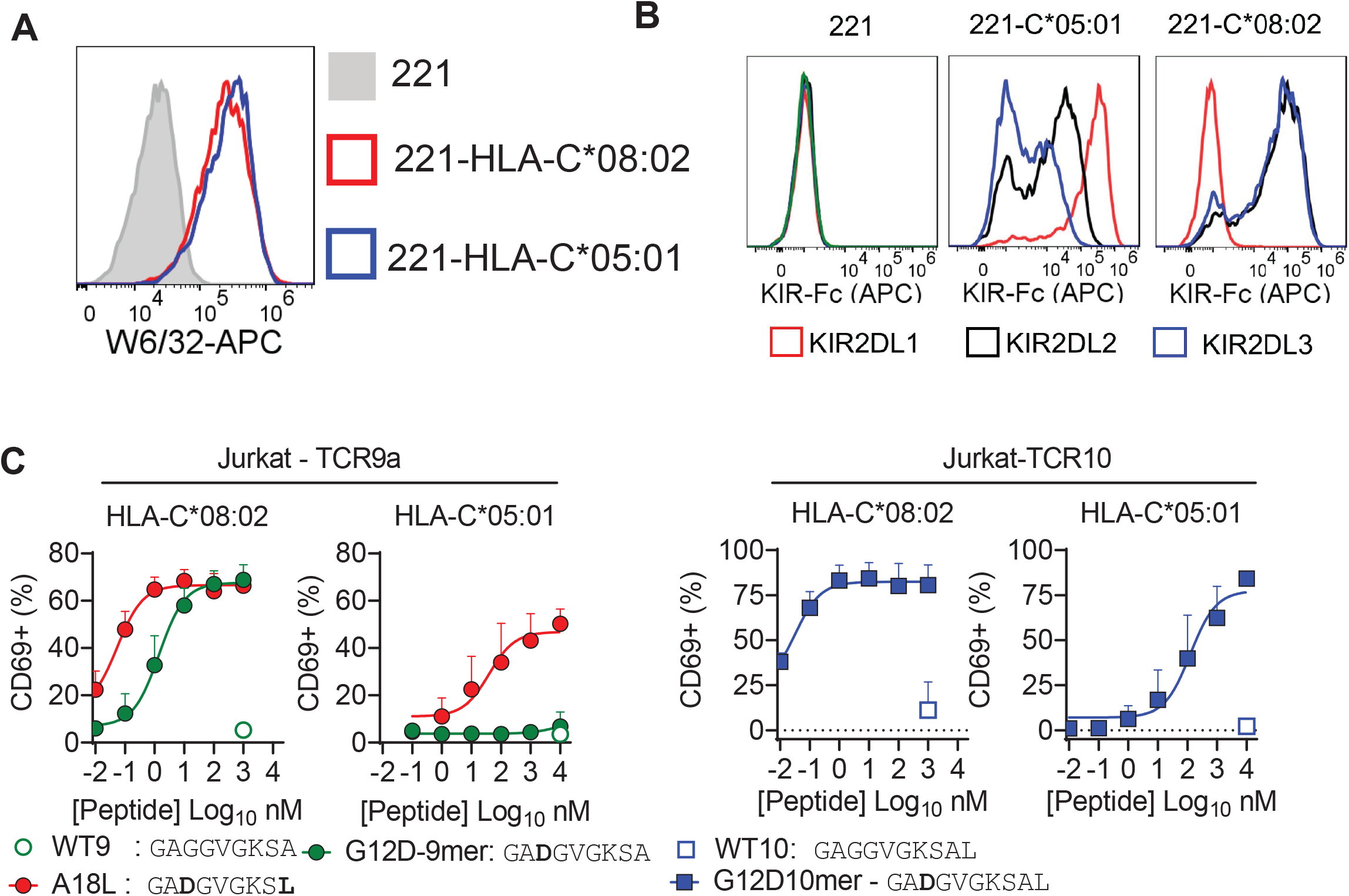
Impact of C1/C2 dimorphism in T cell recognition of HLA-C. **(A)** Expression of HLA-C*08:02 and HLA-C*05:01 on 721.221 (221) cells. **(B)** Recombinant KIR-Ig fusion protein (KIR-Fc) binding to 221, 221-C805:01 and 221-C*08:02. KIR-Fc was conjugated to protein-A APC and used at 3.6 μg/ml. **(C)** Stimulation of Jurkat-TCR9a (*left*) and Jurkat-TCR10 (*right*) with indicated peptides loaded on TAP deficient cells expressing HLA-C*08:02 and HLA-C*05:01. Jurkat T cell activation was measured by CD69 expression by flow cytometry. Means and standard errors from at least three independent experiments are shown.

**SFigure 2.**
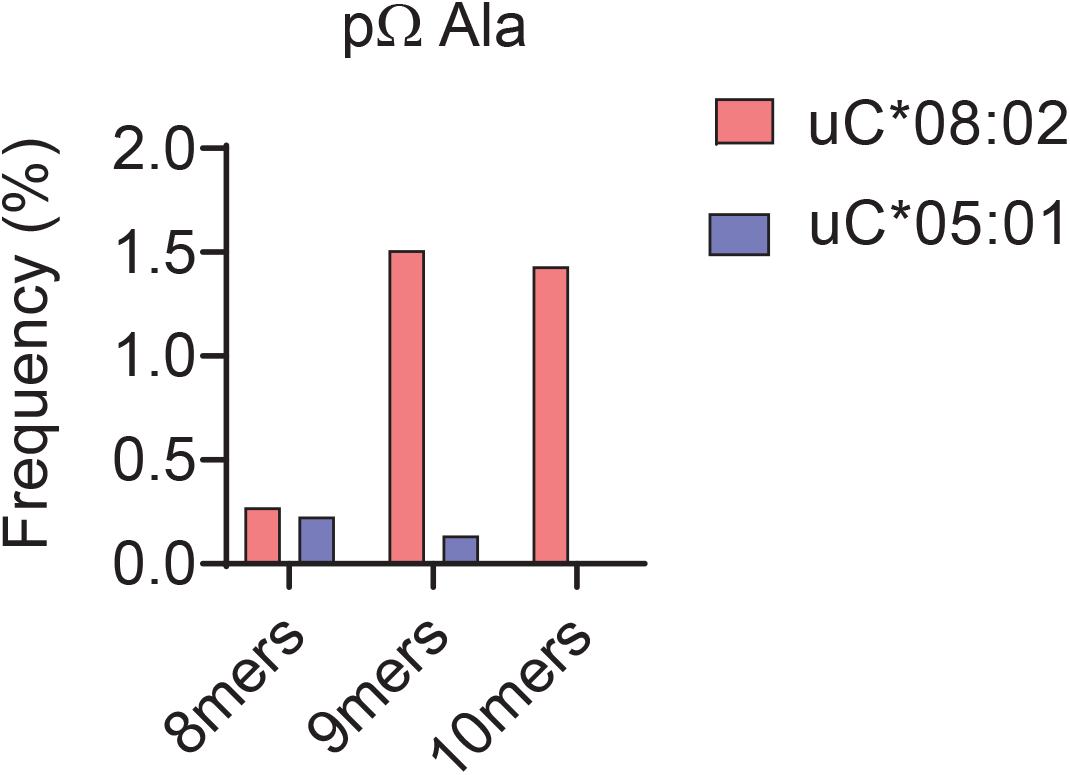
Impact of peptide length on frequency of C-terminal (pΩ) Ala in peptides eluted from C*08:02 and C*05:01. Frequency of peptides with C-terminal Ala in 8mers, 9mers and 10mers eluted from HLA-C*08:02 and HLA-C*05:01.

**SFigure 3.**
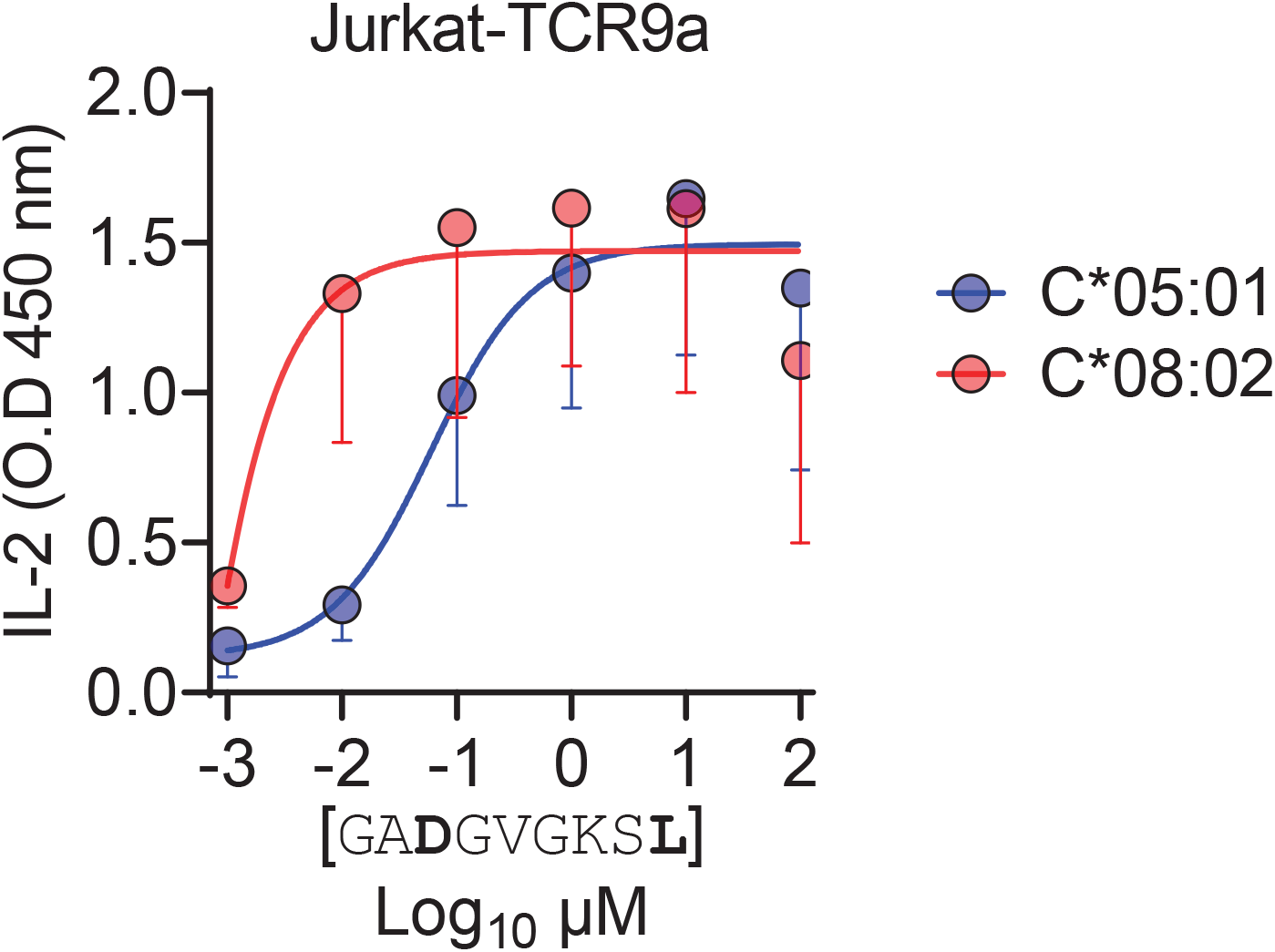
Jurkat-TCR9a is more sensitive to A18L-9mer loaded on C1 C*08:02 than C2 C*05:01. Activation of Jurkat-TCR9a cells by 221-C*08:02 and 221-C*05:01 loaded with A18L-9mer. Means and standard errors of IL-2 concentration in culture supernatant from three independent experiments are shown.

**SFigure 4.**
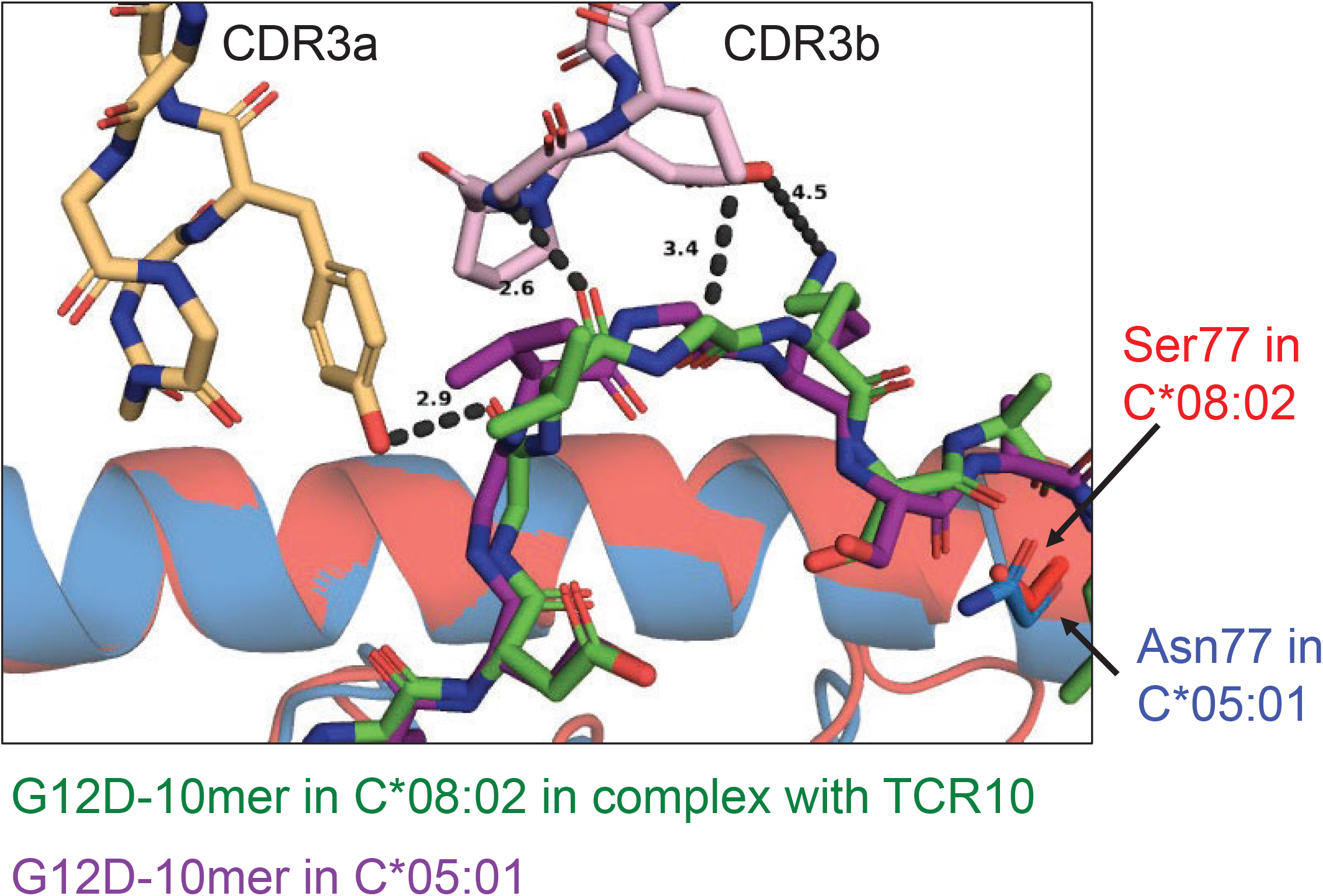
Modelling TCR10 binding to C*05:01-G12D-10mer. Crystal structure of TCR10 in complex with C*08:02-G12D-10mer was used to dock C*05:01-G12D-10mer (PDB:6JTO). Docking was carried out in Pymol.

**SFigure 5.**
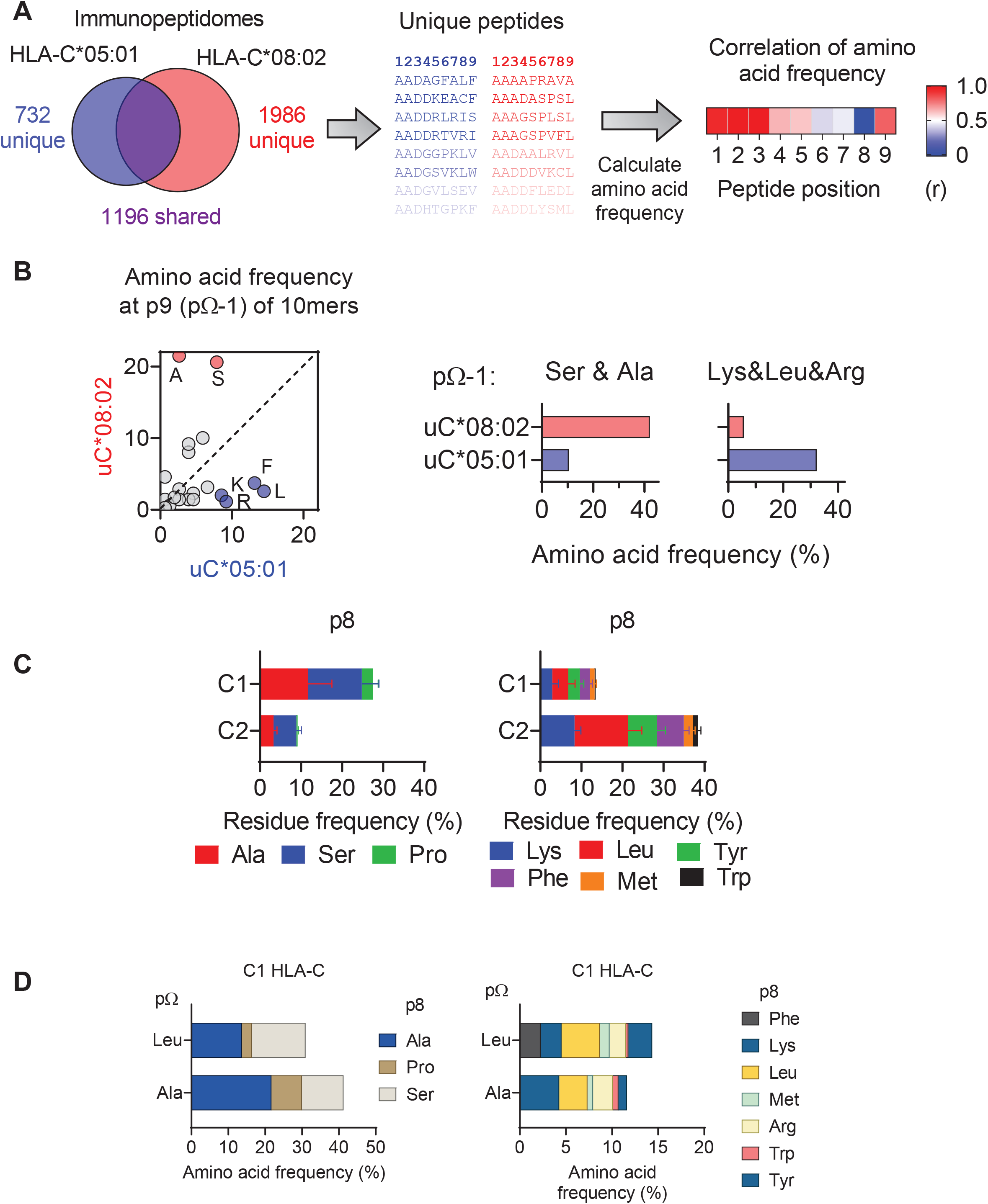
Impact of C1/C2 dimorphism on HLA-C immunopeptidomes. **(A)** Workflow for comparing amino acid frequency of peptides eluted from HLA-C*05:01 and HLA-C*08:02. Amino acid frequencies were compared by Pearson correlation. **(B)** (*left*) Correlation of amino acid frequencies at p9 (pΩ-1) of 10mers eluted from HLA-C*05:01 and HLA-C*08:02. Frequencies of Ser and Ala and Lys, Leu and Arg at pΩ-1 from peptides eluted from HLA-C*08:02 and HLA-C*05:01 (r*ight*). **(C)** Average frequencies of indicated amino acids at p8 (pΩ-1) of 9mers eluted from 14 C1 and 7 C2 HLA-C allotypes. **(D)** Amino acid frequencies at p8 (pΩ-1) of 9mers eluted from C1 allotypes with Ala or Leu at the C-terminus (pΩ).

**SFigure 6.**
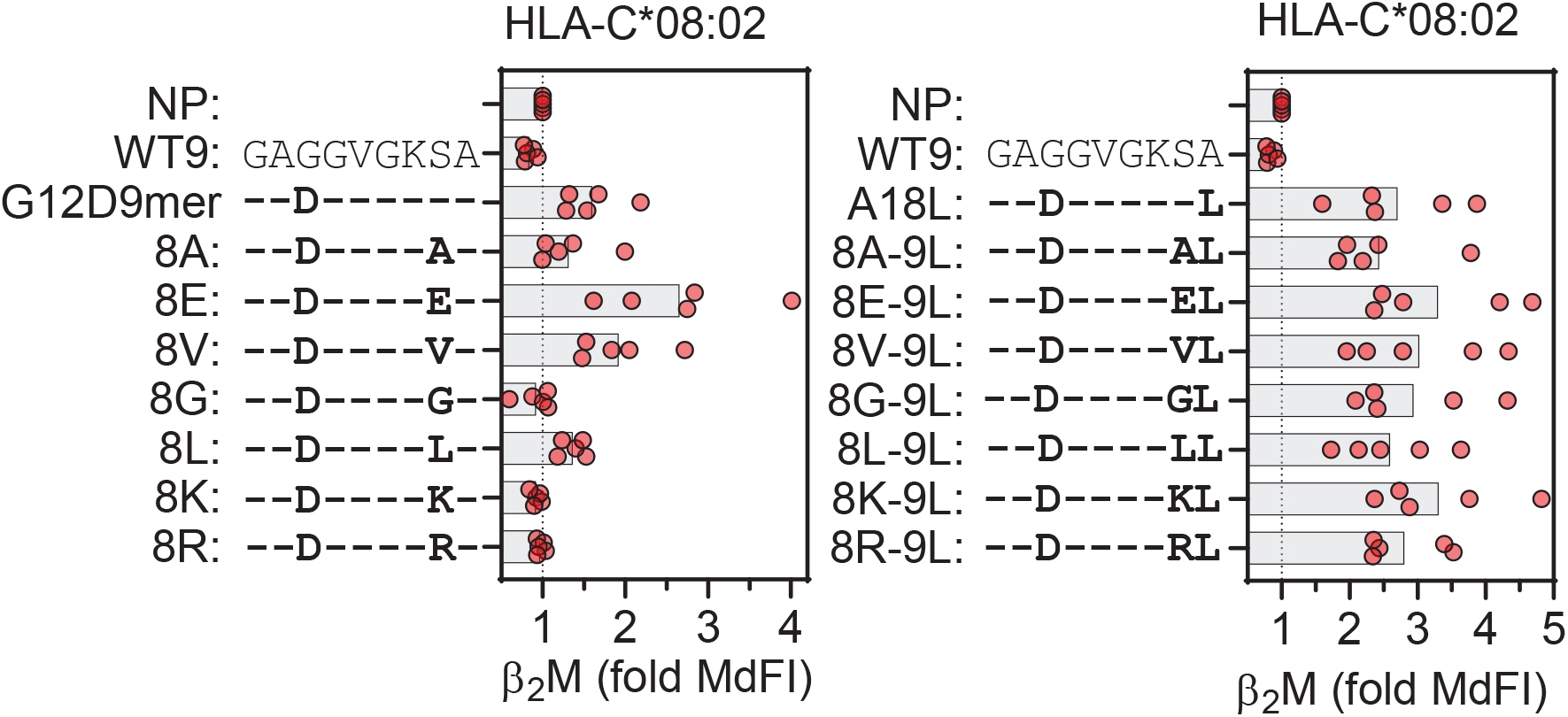
Impact of p8 size on stabilization of HLA-C*08:02. Stabilization of HLA-C on HLA-C*08:02 expressing TAP-deficient cells by G12D-9mer (*left*) and A18L-9mer (*right*) with indicated p8 (pΩ-1) substitutions.

## References

Adams, E.J., and Parham, P. (2001). Species-specific evolution of MHC class I genes in the higher primates. Immunol Rev 183, 41–64.

Adams, P.D., Grosse-Kunstleve, R.W., Hung, L.W., Ioerger, T.R., McCoy, A.J., Moriarty, N.W., Read, R.J., Sacchettini, J.C., Sauter, N.K., and Terwilliger, T.C. (2002). PHENIX: building new software for automated crystallographic structure determination. Acta Crystallogr D Biol Crystallogr 58, 1948–1954.

Addo, M.M., Altfeld, M., Rosenberg, E.S., Eldridge, R.L., Philips, M.N., Habeeb, K., Khatri, A., Brander, C., Robbins, G.K., Mazzara, G.P., et al. (2001). The HIV-1 regulatory proteins Tat and Rev are frequently targeted by cytotoxic T lymphocytes derived from HIV-1-infected individuals. Proc Natl Acad Sci U S A 98, 1781–1786.

Apps, R., Meng, Z., Del Prete, G.Q., Lifson, J.D., Zhou, M., and Carrington, M. (2015). Relative expression levels of the HLA class-I proteins in normal and HIV-infected cells. J Immunol 194, 3594–3600.

Apps, R., Qi, Y., Carlson, J.M., Chen, H., Gao, X., Thomas, R., Yuki, Y., Del Prete, G.Q., Goulder, P., Brumme, Z.L., et al. (2013). Influence of HLA-C expression level on HIV control. Science 340, 87–91.

Bai, P., Zhou, Q., Wei, P., Bai, H., Chan, S.K., Kappler, J.W., Marrack, P., and Yin, L. (2021). Rational discovery of a cancer neoepitope harboring the KRAS G12D driver mutation. Sci China Life Sci.

Biassoni, R., Falco, M., Cambiaggi, A., Costa, P., Verdiani, S., Pende, D., Conte, R., Di Donato, C., Parham, P., and Moretta, L. (1995). Amino acid substitutions can influence the natural killer (NK)-mediated recognition of HLA-C molecules. Role of serine-77 and lysine-80 in the target cell protection from lysis mediated by “group 2” or “group 1” NK clones. J Exp Med 182, 605–609.

Boyington, J.C., Motyka, S.A., Schuck, P., Brooks, A.G., and Sun, P.D. (2000). Crystal structure of an NK cell immunoglobulin-like receptor in complex with its class I MHC ligand. Nature 405, 537–543.

Boyington, J.C., and Sun, P.D. (2002). A structural perspective on MHC class I recognition by killer cell immunoglobulin-like receptors. Mol Immunol 38, 1007–1021.

Chen, L., and Tsai, T.F. (2018). HLA-Cw6 and psoriasis. Br J Dermatol 178, 854–862.

Clements, C.S., Kjer-Nielsen, L., MacDonald, W.A., Brooks, A.G., Purcell, A.W., McCluskey, J., and Rossjohn, J. (2002). The production, purification and crystallization of a soluble heterodimeric form of a highly selected T-cell receptor in its unliganded and liganded state. Acta Crystallogr D Biol Crystallogr 58, 2131–2134.

Collaborative Computational Project, N. (1994). The CCP4 suite: programs for protein crystallography. Acta Crystallogr D Biol Crystallogr 50, 760–763.

Cox, A.D., Fesik, S.W., Kimmelman, A.C., Luo, J., and Der, C.J. (2014). Drugging the undruggable RAS: Mission possible? Nat Rev Drug Discov 13, 828–851.

Debebe, B.J., Boelen, L., Lee, J.C., Investigators, I.P.C., Thio, C.L., Astemborski, J., Kirk, G., Khakoo, S.I., Donfield, S.M., Goedert, J.J., et al. (2020). Identifying the immune interactions underlying HLA class I disease associations. Elife 9.

Dendrou, C.A., Petersen, J., Rossjohn, J., and Fugger, L. (2018). HLA variation and disease. Nat Rev Immunol 18, 325–339.

Devlin, J.R., Alonso, J.A., Ayres, C.M., Keller, G.L.J., Bobisse, S., Vander Kooi, C.W., Coukos, G., Gfeller, D., Harari, A., and Baker, B.M. (2020). Structural dissimilarity from self drives neoepitope escape from immune tolerance. Nat Chem Biol 16, 1269–1276.

Di Marco, M., Schuster, H., Backert, L., Ghosh, M., Rammensee, H.G., and Stevanovic, S. (2017). Unveiling the Peptide Motifs of HLA-C and HLA-G from Naturally Presented Peptides and Generation of Binding Prediction Matrices. J Immunol 199, 2639–2651.

Duru, A.D., Sun, R., Allerbring, E.B., Chadderton, J., Kadri, N., Han, X., Peqini, K., Uchtenhagen, H., Madhurantakam, C., Pellegrino, S., et al. (2020). Tuning antiviral CD8 T-cell response via proline-altered peptide ligand vaccination. PLoS Pathog 16, e1008244.

Emsley, P., and Cowtan, K. (2004). Coot: model-building tools for molecular graphics. Acta Crystallogr D Biol Crystallogr 60, 2126–2132.

Falk, K., Rotzschke, O., Stevanovic, S., Jung, G., and Rammensee, H.G. (1991). Allele-specific motifs revealed by sequencing of self-peptides eluted from MHC molecules. Nature 351, 290–296.

Fan, Q.R., Long, E.O., and Wiley, D.C. (2001). Crystal structure of the human natural killer cell inhibitory receptor KIR2DL1-HLA-Cw4 complex. Nat Immunol 2, 452–460.

Garboczi, D.N., Hung, D.T., and Wiley, D.C. (1992). HLA-A2-peptide complexes: refolding and crystallization of molecules expressed in Escherichia coli and complexed with single antigenic peptides. Proc Natl Acad Sci U S A 89, 3429–3433.

Hiby, S.E., Walker, J.J., O’Shaughnessy K M., Redman, C.W., Carrington, M., Trowsdale, J., and Moffett, A. (2004). Combinations of maternal KIR and fetal HLA-C genes influence the risk of preeclampsia and reproductive success. J Exp Med 200, 957–965.

Hilton, H.G., and Parham, P. (2017). Missing or altered self: human NK cell receptors that recognize HLA-C. Immunogenetics 69, 567–579.

Illing, P.T., Pymm, P., Croft, N.P., Hilton, H.G., Jojic, V., Han, A.S., Mendoza, J.L., Mifsud, N.A., Dudek, N.L., McCluskey, J., et al. (2018). HLA-B57 micropolymorphism defines the sequence and conformational breadth of the immunopeptidome. Nat Commun 9, 4693.

Illing, P.T., Vivian, J.P., Dudek, N.L., Kostenko, L., Chen, Z., Bharadwaj, M., Miles, J.J., Kjer-Nielsen, L., Gras, S., Williamson, N.A., et al. (2012). Immune self-reactivity triggered by drug-modified HLA-peptide repertoire. Nature 486, 554–558.

Istrail, S., Florea, L., Halldorsson, B.V., Kohlbacher, O., Schwartz, R.S., Yap, V.B., Yewdell, J.W., and Hoffman, S.L. (2004). Comparative immunopeptidomics of humans and their pathogens. Proc Natl Acad Sci U S A 101, 13268–13272.

Khakoo, S.I., Thio, C.L., Martin, M.P., Brooks, C.R., Gao, X., Astemborski, J., Cheng, J., Goedert, J.J., Vlahov, D., Hilgartner, M., et al. (2004). HLA and NK cell inhibitory receptor genes in resolving hepatitis C virus infection. Science 305, 872–874.

King, A., Burrows, T.D., Hiby, S.E., Bowen, J.M., Joseph, S., Verma, S., Lim, P.B., Gardner, L., Le Bouteiller, P., Ziegler, A., et al. (2000). Surface expression of HLA-C antigen by human extravillous trophoblast. Placenta 21, 376–387.

Kulkarni, S., Martin, M.P., and Carrington, M. (2008). The Yin and Yang of HLA and KIR in human disease. Semin Immunol 20, 343–352.

Lande, R., Botti, E., Jandus, C., Dojcinovic, D., Fanelli, G., Conrad, C., Chamilos, G., Feldmeyer, L., Marinari, B., Chon, S., et al. (2014). The antimicrobial peptide LL37 is a T-cell autoantigen in psoriasis. Nat Commun 5, 5621.

Levin, N., Paria, B.C., Vale, N.R., Yossef, R., Lowery, F.J., Parkhurst, M.R., Yu, Z., Florentin, M., Cafri, G., Gartner, J.J., et al. (2021). Identification and Validation of T-cell Receptors Targeting RAS Hotspot Mutations in Human Cancers for Use in Cell-based Immunotherapy. Clin Cancer Res 27, 5084–5095.

Littera, R., Chessa, L., Onali, S., Figorilli, F., Lai, S., Secci, L., La Nasa, G., Caocci, G., Arras, M., Melis, M., et al. (2016). Exploring the Role of Killer Cell Immunoglobulin-Like Receptors and Their HLA Class I Ligands in Autoimmune Hepatitis. PLoS One 11, e0146086.

Lu, Y.C., Yao, X., Crystal, J.S., Li, Y.F., El-Gamil, M., Gross, C., Davis, L., Dudley, M.E., Yang, J.C., Samuels, Y., et al. (2014). Efficient identification of mutated cancer antigens recognized by T cells associated with durable tumor regressions. Clin Cancer Res 20, 3401–3410.

Madden, D.R. (1995). The three-dimensional structure of peptide-MHC complexes. Annu Rev Immunol 13, 587–622.

McCoy, A.J., Grosse-Kunstleve, R.W., Adams, P.D., Winn, M.D., Storoni, L.C., and Read, R.J. (2007). Phaser crystallographic software. J Appl Crystallogr 40, 658–674.

McCutcheon, J.A., Gumperz, J., Smith, K.D., Lutz, C.T., and Parham, P. (1995). Low HLA-C expression at cell surfaces correlates with increased turnover of heavy chain mRNA. J Exp Med 181, 2085–2095.

Moesta, A.K., Norman, P.J., Yawata, M., Yawata, N., Gleimer, M., and Parham, P. (2008). Synergistic polymorphism at two positions distal to the ligand-binding site makes KIR2DL2 a stronger receptor for HLA-C than KIR2DL3. J Immunol 180, 3969–3979.

Murata, K., Nakatsugawa, M., Rahman, M.A., Nguyen, L.T., Millar, D.G., Mulder, D.T., Sugata, K., Saijo, H., Matsunaga, Y., Kagoya, Y., et al. (2020). Landscape mapping of shared antigenic epitopes and their cognate TCRs of tumor-infiltrating T lymphocytes in melanoma. Elife 9.

Otwinowski, Z., and Minor, W. (1997). Processing of X-ray diffraction data collected in oscillation mode. Methods Enzymol 276, 307–326.

Parham, P., and Moffett, A. (2013). Variable NK cell receptors and their MHC class I ligands in immunity, reproduction and human evolution. Nat Rev Immunol 13, 133–144.

Parham, P., Norman, P.J., Abi-Rached, L., and Guethlein, L.A. (2012). Human-specific evolution of killer cell immunoglobulin-like receptor recognition of major histocompatibility complex class I molecules. Philos Trans R Soc Lond B Biol Sci 367, 800–811.

Pearlman, A.H., Hwang, M.S., Konig, M.F., Hsiue, E.H.-C., Douglass, J., DiNapoli, S.R., Mog, B.J., Bettegowda, C., Pardoll, D.M., Gabelli, S.B., et al. (2021). Targeting public neoantigens for cancer immunotherapy. Nature Cancer 2, 487–497.

Rajagopalan, S., and Long, E.O. (2005). Understanding how combinations of HLA and KIR genes influence disease. J Exp Med 201, 1025–1029.

Robinson, J., Guethlein, L.A., Cereb, N., Yang, S.Y., Norman, P.J., Marsh, S.G.E., and Parham, P. (2017). Distinguishing functional polymorphism from random variation in the sequences of >10,000 HLA-A, -B and -C alleles. PLoS Genet 13, e1006862.

Rossjohn, J., Gras, S., Miles, J.J., Turner, S.J., Godfrey, D.I., and McCluskey, J. (2015). T cell antigen receptor recognition of antigen-presenting molecules. Annu Rev Immunol 33, 169–200.

Sarkizova, S., Klaeger, S., Le, P.M., Li, L.W., Oliveira, G., Keshishian, H., Hartigan, C.R., Zhang, W., Braun, D.A., Ligon, K.L., et al. (2020). A large peptidome dataset improves HLA class I epitope prediction across most of the human population. Nat Biotechnol 38, 199–209.

Saunders, P.M., MacLachlan, B.J., Pymm, P., Illing, P.T., Deng, Y., Wong, S.C., Oates, C.V.L., Purcell, A.W., Rossjohn, J., Vivian, J.P., et al. (2020). The molecular basis of how buried human leukocyte antigen polymorphism modulates natural killer cell function. Proc Natl Acad Sci U S A 117, 11636–11647.

Saunders, P.M., Vivian, J.P., O’Connor, G.M., Sullivan, L.C., Pymm, P., Rossjohn, J., and Brooks, A.G. (2015). A bird’s eye view of NK cell receptor interactions with their MHC class I ligands. Immunol Rev 267, 148–166.

Sim, M.J., Malaker, S.A., Khan, A., Stowell, J.M., Shabanowitz, J., Peterson, M.E., Rajagopalan, S., Hunt, D.F., Altmann, D.M., Long, E.O., et al. (2017). Canonical and Cross-reactive Binding of NK Cell Inhibitory Receptors to HLA-C Allotypes Is Dictated by Peptides Bound to HLA-C. Front Immunol 8, 193.

Sim, M.J.W., Lu, J., Spencer, M., Hopkins, F., Tran, E., Rosenberg, S.A., Long, E.O., and Sun, P.D. (2020). High-affinity oligoclonal TCRs define effective adoptive T cell therapy targeting mutant KRAS-G12D. Proc Natl Acad Sci U S A.

Sim, M.J.W., Rajagopalan, S., Altmann, D.M., Boyton, R.J., Sun, P.D., and Long, E.O. (2019). Human NK cell receptor KIR2DS4 detects a conserved bacterial epitope presented by HLA-C. Proc Natl Acad Sci U S A 116, 12964–12973.

Stephen, A.G., Esposito, D., Bagni, R.K., and McCormick, F. (2014). Dragging ras back in the ring. Cancer Cell 25, 272–281.

Stewart-Jones, G.B., Simpson, P., van der Merwe, P.A., Easterbrook, P., McMichael, A.J., Rowland-Jones, S.L., Jones, E.Y., and Gillespie, G.M. (2012). Structural features underlying T-cell receptor sensitivity to concealed MHC class I micropolymorphisms. Proc Natl Acad Sci U S A 109, E3483–3492.

Tikhonova, A.N., Van Laethem, F., Hanada, K., Lu, J., Pobezinsky, L.A., Hong, C., Guinter, T.I., Jeurling, S.K., Bernhardt, G., Park, J.H., et al. (2012). alphabeta T cell receptors that do not undergo major histocompatibility complex-specific thymic selection possess antibody-like recognition specificities. Immunity 36, 79–91.

Tran, E., Ahmadzadeh, M., Lu, Y.C., Gros, A., Turcotte, S., Robbins, P.F., Gartner, J.J., Zheng, Z., Li, Y.F., Ray, S., et al. (2015). Immunogenicity of somatic mutations in human gastrointestinal cancers. Science 350, 1387–1390.

Tran, E., Robbins, P.F., Lu, Y.C., Prickett, T.D., Gartner, J.J., Jia, L., Pasetto, A., Zheng, Z., Ray, S., Groh, E.M., et al. (2016). T-Cell Transfer Therapy Targeting Mutant KRAS in Cancer. N Engl J Med 375, 2255–2262.

Tynan, F.E., Elhassen, D., Purcell, A.W., Burrows, J.M., Borg, N.A., Miles, J.J., Williamson, N.A., Green, K.J., Tellam, J., Kjer-Nielsen, L., et al. (2005). The immunogenicity of a viral cytotoxic T cell epitope is controlled by its MHC-bound conformation. J Exp Med 202, 1249–1260.

Venstrom, J.M., Pittari, G., Gooley, T.A., Chewning, J.H., Spellman, S., Haagenson, M., Gallagher, M.M., Malkki, M., Petersdorf, E., Dupont, B., et al. (2012). HLA-C-dependent prevention of leukemia relapse by donor activating KIR2DS1. N Engl J Med 367, 805–816.

